# *TRIM71* deficiency causes germ cell loss during mouse embryogenesis and promotes human male infertility

**DOI:** 10.1101/2021.02.01.429172

**Authors:** Lucia A. Torres-Fernández, Jana Emich, Yasmine Port, Sibylle Mitschka, Marius Wöste, Simon Schneider, Daniela Fietz, Manon S. Oud, Sara Di Persio, Nina Neuhaus, Sabine Kliesch, Michael Hölzel, Hubert Schorle, Corinna Friedrich, Frank Tüttelmann, Waldemar Kolanus

**Author notes:** Equal contribution (Lucia A. Torres-Fernández, Jana Emich, Yasmine Port, Frank Tüttelmann, Waldemar Kolanus).

## Abstract

Mutations affecting the germline can result in infertility or the generation of germ cell tumors (GCT), highlighting the need to identify and characterize the genes controlling the complex molecular network orchestrating germ cell development. TRIM71 is a stem cell-specific factor essential for embryogenesis, and its expression has been reported in GCT and adult mouse testes. To investigate the role of TRIM71 in mammalian germ cell embryonic development, we generated a germline-specific conditional *Trim71* knockout mouse (cKO) using the early primordial germ cell (PGC) marker *Nanos3* as a Cre-recombinase driver. cKO mice are infertile, with male mice displaying a Sertoli cell-only (SCO) phenotype, which in humans is defined as a specific subtype of non-obstructive azoospermia characterized by the absence of developing germ cells in the testes’ seminiferous tubules. Infertility originates during embryogenesis, as the SCO phenotype was already apparent in neonatal mice. The *in vitro* differentiation of mouse embryonic stem cells (ESCs) into PGC-like cells (PGCLCs) revealed reduced numbers of PGCLCs in *Trim71*-deficient cells. Furthermore, *in vitro* growth competition assays with wild type and CRISPR/Cas9-generated *TRIM71* mutant NCCIT cells, a human GCT-derived cell line which we used as a surrogate model for proliferating PGCs, showed that TRIM71 promotes NCCIT cell proliferation and survival. Our data collectively suggest that germ cell loss in cKO mice results from combined defects during the specification and maintenance of PGCs prior to their sex determination in the genital ridges. Last, via exome sequencing analysis, we identified several *TRIM71* variants in a cohort of infertile men, including a loss-of-function variant in a patient with SCO phenotype. Our work reveals for the first time an association of *TRIM71* variants with human male infertility, and uncovers further developmental roles for TRIM71 in the generation and maintenance of germ cells during mouse embryogenesis.

## Introduction

Despite the recent insights into the regulation of mammalian germ cell development and the advances in assisted reproductive technologies, infertility remains a common problem in modern society affecting ∼15% of couples in industrialized countries^1^. Genetic defects during germ cell development can drive infertility and increase susceptibility to germ cell tumors (GCT)^2–4^.

Primordial germ cells (PGCs) are the first established germ cell population during embryonic development. In mice, PGC specification is initiated in the post-implantation epiblast at embryonic day (E) 6.25 in response to BMP4 signaling from the extra-embryonic ectoderm^5^. BMP4 activates the expression of *Prdm14*^6^ and *Prdm1/Blimp1*^7^. BLIMP1 then activates the expression of *Tfap2c*^8^, and together BLIMP1, TFAP2C and PRDM14 establish the transcriptional program required for PGC specification^9^ which is completed at E7.5. At this time point, about 40 cells in the proximal epiblast express early PGC-specific markers such as *Nanos3*^10,11^. PGCs then migrate to the genital ridges (developing gonads) while slowly proliferating^12,13^. At E10.5, around 1000 PGCs colonize the genital ridges, upregulate late PGC markers such as *Ddx4/Vasa*^14,15^ and *Gcna1*^16^, and undergo significant proliferation before initiating sex determination^17,18^. Importantly, PGCs that fail to reach the genital ridges or to further differentiate into gametes are the origin of GCT, which often affect children and young adults^2–4,19^. Therefore, unravelling the processes controlling germline development will not only contribute to our understanding of infertility, but may also offer unique opportunities for the treatment of GCT.

Tripartite Motif Containing 71 (TRIM71) belongs to the TRIM-NHL family and is a stem cell-specific protein^20^ which can act both as an E3 ubiquitin ligase^21–24^ and an mRNA repressor^25–29^. TRIM71 function is essential for embryogenesis and its expression is mostly restricted to early proliferative developmental stages, being downregulated in the course of differentiation^30–33^. However, a postnatal *Trim71* expression has also been observed in adult mouse testes^33,34^ as well as in several GCT-derived cell lines^25,29,33^, and a recent study has reported a postnatal function for TRIM71 in adult mouse spermatogenesis^34^. Furthermore, RNA sequencing of *Trim71*-deficient mouse embryonic stem cells (ESCs) revealed a decreased expression of genes associated with reproductive processes^28^, suggesting an early developmental role for TRIM71 in the germline.

To elucidate the role of TRIM71 in fertility and mammalian embryonic germ cell development, we generated a mouse model with an early germline-specific depletion of *Trim71* driven by the *Nanos3*-Cre promoter. Additionally, we employed an *in vitro* approach for the differentiation of wild type and *Trim71*-deficient ESCs into PGCs in order to study the role of TRIM71 during PGC specification. Furthermore, we generated *TRIM71* mutant NCCIT cells via CRISPAR/Cas9 to investigate the role of TRIM71 in the proliferation of GCT-derived cells. Last, we used exome sequencing data from infertile men and developed a novel software tool named Haystack to search for novel genetic causes of infertility. Our work thereby identifies *TRIM71* as a novel gene associated with human male infertility, and uncovers further developmental roles for TRIM71 in the generation and maintenance of germ cells during mouse embryogenesis.

## Results

### *Trim71* is expressed in spermatogonial stem cells and is essential for mouse fertility

Mice carrying a *Trim71* homozygous deletion (KO, *Trim71*^*-/-*^) die during embryonic development (Suppl. Fig. 1A)^31,32^. In contrast, *Trim71* heterozygous mice (HET, *Trim71*^*fl/-*^) are viable and fertile, although significantly smaller in length and weight than wild type (WT, *Trim71*^*fl/fl*^) littermates (Suppl. Fig. 1B-C). We detected *Trim71* expression in the testes of wild type and heterozygous adult mice – but not in other organs such as heart or kidney – with *Trim71* mRNA levels decreased in the testis of heterozygous mice (Fig. 1A). We then measured the weight of several organs relative to the total body weight, and observed that testes – but neither heart nor kidney – were significantly smaller in heterozygous males compared to wild type males (Fig. 1B). Accordingly, sperm counts were significantly reduced in adult heterozygous males (Fig. 1C). These data suggest that *Trim71* expression levels are important for male gonad development and reveal a haploinsufficiency of *Trim71* in mice.

**Figure 1.**
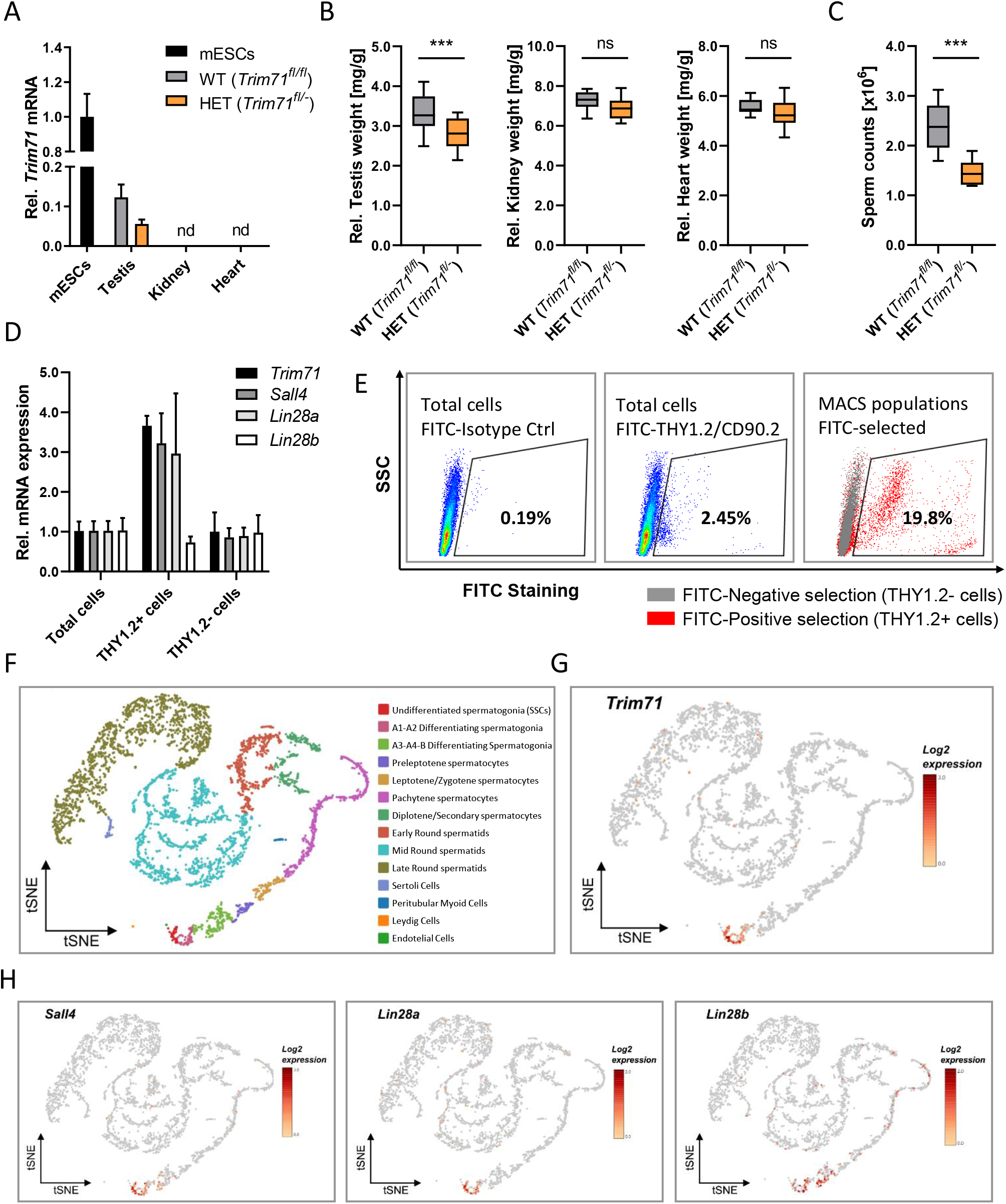
*Trim71* is expressed in spermatogonial stem cells (SSCs) of adult fertile mice. **A)** qRT-PCR of *Trim71* relative to *Hprt* housekeeping gene in testis, kidney and heart of wild type (WT, *Trim71*^*fl/fl*^) and *Trim71* heterozygous (HET, *Trim71*^*fl/-*^) adult male mice (10-14 weeks old), normalized to *Trim71* relative levels in wild type murine ESCs (nd = not detected). Error bars represent SD (n=3). **B)** Organ weight [mg] relative to total body weight [g] of testis, kidney and heart of wild type (WT, *Trim71*^*fl/fl*^) and *Trim71* heterozygous (HET, *Trim71*^*fl/*-^) male adult mice (10-14 weeks old). Graphs represent Tukey plots (n=14-16). ***P-value < 0.005, ns = non-significant (unpaired Student’s t-test). **C)** Epididymal sperm counts in wild type (WT, *Trim71*^*fl/fl*^) and *Trim71* heterozygous (HET, *Trim71*^*fl/-*^) male adult mice (10-14 weeks old). ***P-value < 0.005 (unpaired Student’s t-test). **D)** qRT-PCR of *Trim71*, the SSC markers *Sall4* and *Lin28a*, and the differentiating spermatogonia marker *Lin28b*, relative to *Hprt* housekeeping gene in testes cell suspension before (total cells) and after THY1.2 MACS (THY1.2+/- cells). Error bars represent SD (n=3). **E)** Representative flow cytometry scatter plots of testes cell suspension before (total cells) and after THY1.2/CD90.2 MACS (Thy1.2+/- cells). **F)** tSNE plot (t-distributed stochastic neighbor embedding) of single-cell transcriptome data (scRNA-seq) from mouse testes as published by Hermann et al., 2018. Each dot represents a single cell and is colored according to its cluster identity as indicated on the figure key. G) Expression pattern of *Trim71* and H) *Sall4, Lin28a* and *Lin28b* projected on the tSNE plot of the mouse scRNA-seq dataset. Red indicates high expression and gray indicates low or no expression, as shown by the figure key. See also Suppl. Fig. 1.

Since TRIM71 is a stem cell-specific protein, we hypothesized that its expression in mice testes is present in undifferentiated spermatogonia, also known as spermatogonial stem cells (SSCs). To validate this hypothesis, we used wild type testes cell suspensions to enrich SSCs (THY1.2+) via magnetic-activated cell sorting (MACS)^35^. We found an increase of *Trim71* mRNA as well as the mRNAs of the known SSC-specific markers *Sall4*^36^ and *Lin28a*^37^ – but not the differentiating spermatogonia marker *Lin28b*^38^ – in the THY1.2+ enriched cell population (Fig. 1D-E). This result indicates that *Trim71* expression in adult mouse testes is enriched in SSCs. Indeed, single-cell RNA sequencing of mouse adult testicular tissue confirms a restricted expression of *Trim71* in SSCs (Fig. 1F-H)^39^.

To study the function of *Trim71* in male gonad development, a germline-specific *Trim71* conditional knockout mouse (cKO) was generated with the Cre recombinase expressed under the promoter of the early PGC marker *Nanos3* (Suppl. Fig. 2A). For a first functional evaluation, adult cKO mice (*Trim71*^*fl/-*^; *Nanos3*^*Cre/+*^) were crossed with wild type mice. *Trim71* deficiency in the germline resulted in infertility in both sexes, as neither males nor females were able to produce offspring (Suppl. Fig. 2B).

### Germline-specific *Trim71* cKO male mice display a Sertoli cell-only (SCO)-like phenotype

Macroscopic analysis of cKO male and female reproductive organs revealed a dramatic reduction in testis and ovary size, respectively (Fig. 2A-C and Suppl. Fig. 2C). Hematoxylin and eosin (H&E) staining of adult male testis cross-sections showed that most seminiferous tubules in cKO testis were lacking signs of spermatogenesis (Fig. 2D). Furthermore, immunofluorescence staining of germ cells (GCNA1^40,41^) and Sertoli cells (WT1^42^) showed a dramatic reduction of seminiferous tubules containing GCNA+ cells in cKO mice (Fig. 2E-F). Accordingly, the expression of SSC-specific markers *Oct4, Sall4* and *Lin28a*, as well as the differentiating spermatogonia marker *Lin28b*, were dramatically reduced in whole testis RNA of cKO mice (Fig. 2G), representing the deficit of germ cells in these mice.

**Figure 2.**
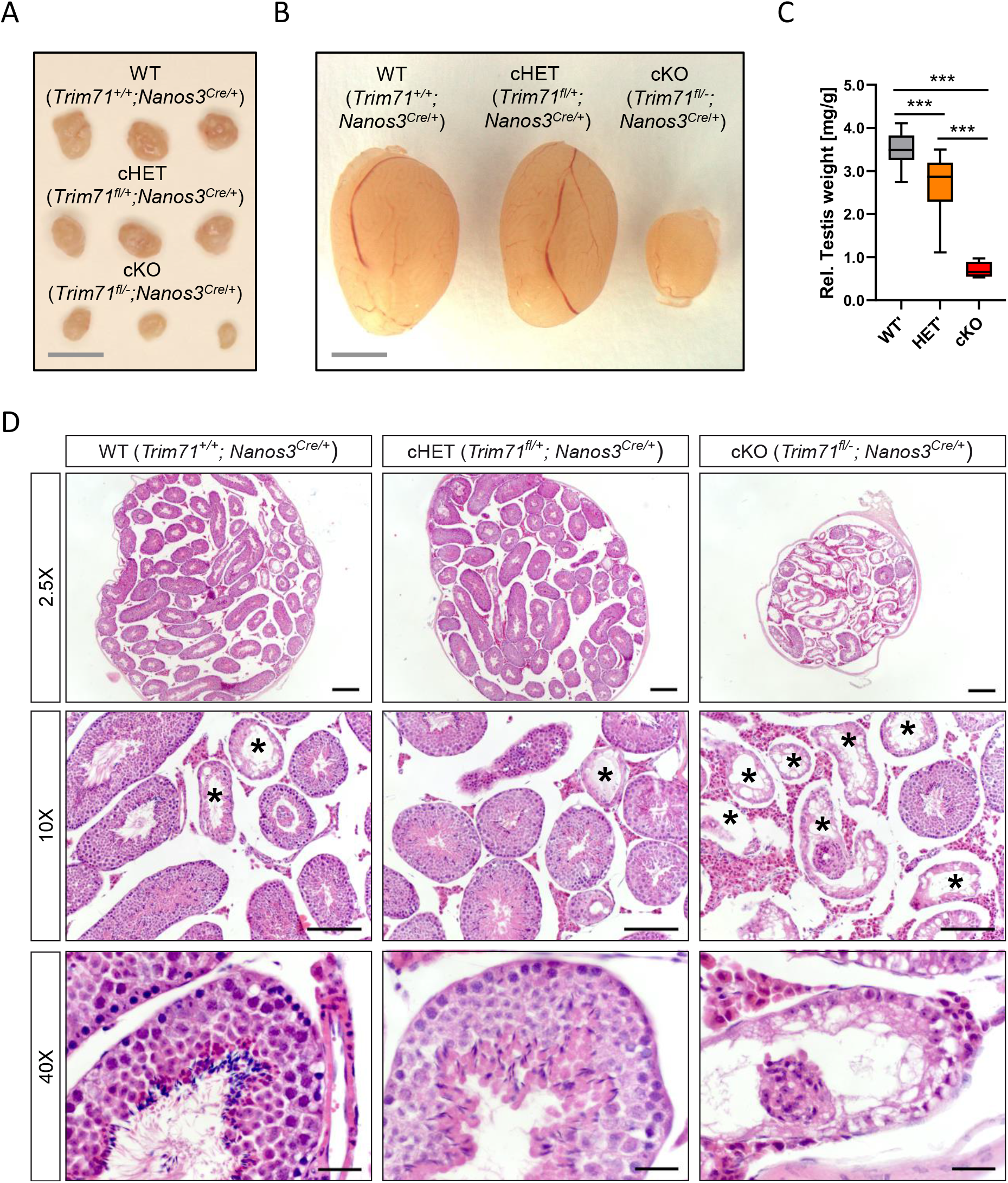
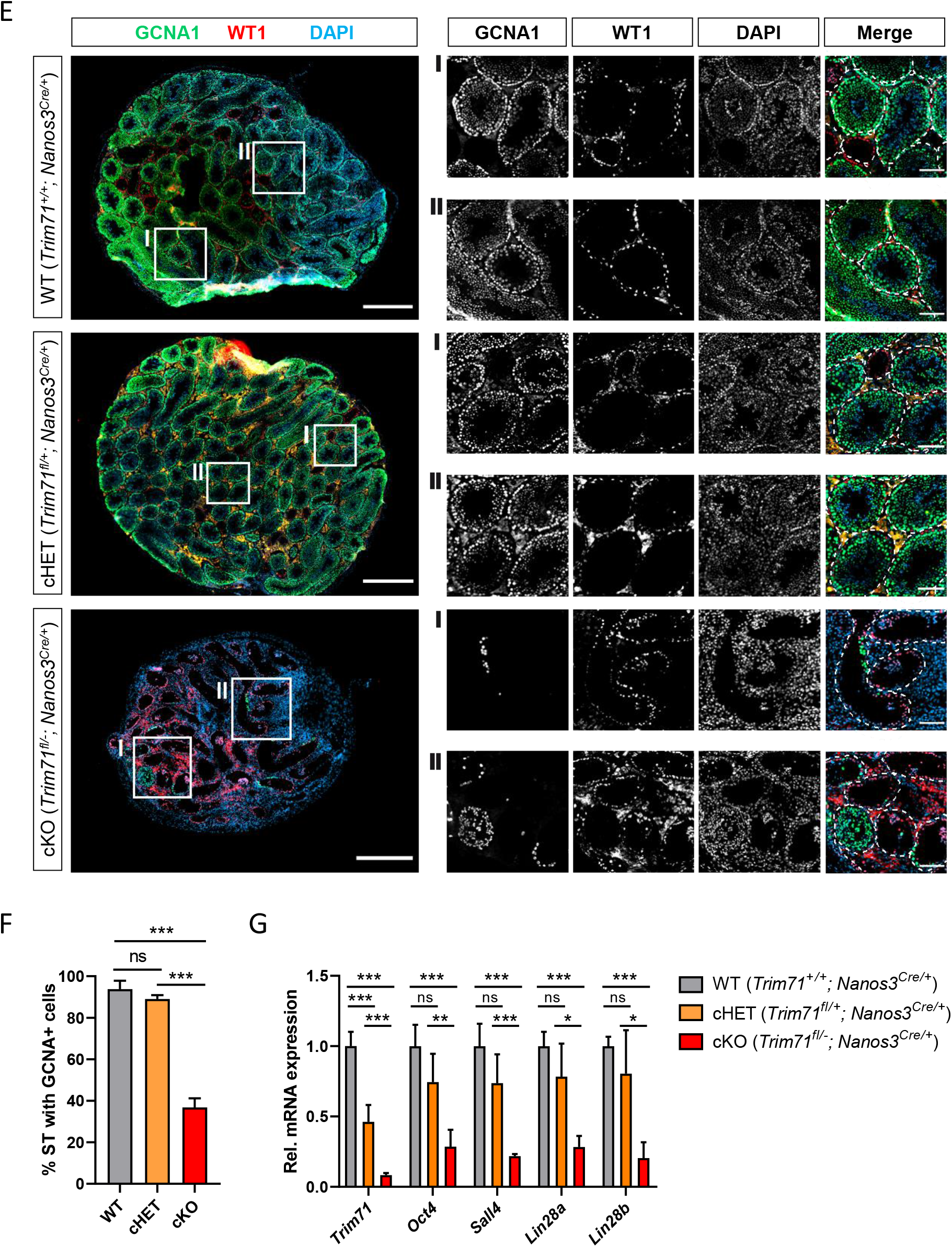
Germline-specific *Trim71* cKO male mice present a Sertoli cell-only (SCO) phenotype. **A)** Representative images of ovaries and **B)** testes of adult wild type (WT, *Trim71*^+/+^; *Nanos3*^Cre/+^), germline-specific *Trim71* heterozygous (cHET, *Trim71*^fl/+^; *Nanos3*^Cre/+^) and germline-specific *Trim71* knockout (cKO, *Trim71*^fl/-^; *Nanos3*^Cre/+^) mice. Scale bars represent 2 mm. **C)** Testis weight [mg] relative to total body weight [g] of wild type (WT’), *Trim71* heterozygous (HET’) and germline-specific *Trim71* knockout (cKO) mice, summarized from Suppl. Fig. 2C, joining several *Nanos3* genotypes per *Trim71* genotype. Error bars represent SD (n=8-29). *** P-value < 0.005 (one-way ANOVA, Tukey’s test). **D)** Representative H&E stainings on paraffin testes cross-sections from adult wild type (WT, *Trim71*^+/+^; *Nanos3*^Cre/+^), germline-specific *Trim71* heterozygous (cHET, *Trim71*^fl/+^; *Nanos3*^Cre/+^) and germline-specific *Trim71* knockout (cKO, *Trim71*^fl/-^; *Nanos3*^Cre/+^) mice. Seminiferous tubules with a defective morphology are marked with an asterisk (*) in 10x magnification images. Scale bars represent 200 μm, 100 μm and 20 μm in 2.5x, 10x and 40x magnifications, respectively. **E)** Representative immunofluorescence stainings on testes cryosections from adult wild type (WT, *Trim71*^+/+^; *Nanos3*^Cre/+^), germline-specific *Trim71* heterozygous (cHET, *Trim71*^fl/+^; *Nanos3*^Cre/+^) and germline-specific *Trim71* knockout (cKO, *Trim71*^fl/-^; *Nanos3*^Cre/+^) mice. Images show co-staining with GCNA1 (germ cells), WT1 (Sertoli cells) and DAPI (nuclei). For each genotype two regions – indicated as I and II – are depicted in higher magnification. Scale bars represent 500 μm for complete testes cross-sections and 100 μm for the magnification images. **F)** Quantification of seminiferous tubules (ST) containing GCNA+ cells per testis cross-section from E, depicted as percentages. Error bars represent SD (n=3). ***P-value < 0.005, ns = non-significant (unpaired Student’s t-test). **G)** qRT-PCR of *Trim71*, the SSC markers *Sall4, Lin28a* and *Oct4*, and the differentiating spermatogonia marker *Lin28b*, relative to *Hprt* housekeeping gene in whole testis RNA of wild type (WT, *Trim71*^+/+^; *Nanos3*^Cre/+^), germline-specific *Trim71* heterozygous (cHET, *Trim71*^fl/+^; *Nanos3*^Cre/+^) and germline-specific *Trim71* knockout (cKO, *Trim71*^fl/-^; *Nanos3*^Cre/+^) male adult mice. Error bars represent SD (n=3-6). ***P-value < 0.005; **P-value < 0.001; *P-value < 0.05; ns = non-significant (unpaired Student’s t-test). See also Suppl. Fig. 2.

This phenotype is reminiscent of a human condition known as Sertoli cell-only (SCO) phenotype, which is a specific type of non-obstructive azoospermia characterized by a total or substantial absence of germ cells in the seminiferous tubules, and whose causes remain mostly unknown^43–45^. Our results here suggest that *Trim71* cKO male mice display an SCO-like phenotype with most seminiferous tubules lacking germ cells.

### Infertility in germline-specific *Trim71* cKO male mice has an embryonic origin

As *Nanos3* expression in mice embryos is detected as early as E7.5 during PGC specification^11^, and sex determination of PGCs does not occur until E10.5^17,18^, the infertility observed in both male and female *Trim71* cKO mice may reflect a function of TRIM71 in an early stage of germ cell development. In order to determine whether the deficit of germ cells derives from defects during embryonic development, we conducted H&E staining and GCNA1-WT1 immunostaining on testis cross-sections from neonatal (P0.5) male mice (Fig. 3A-B). Indeed, the SCO-like phenotype previously observed in our adult cKO male mice was already apparent in neonatal cKO mice with a reduction in gonocyte-containing seminiferous tubules (Fig. 3B-C). A recent study has shown that postnatal *Trim71* expression in male mouse is important for the successful establishment and differentiation of the adult SSC pool^34^. Our results here additionally show that *Trim71* expression during embryonic development is required for the generation and/or maintenance of the PGC/gonocyte pool before birth.

**Figure 3.**
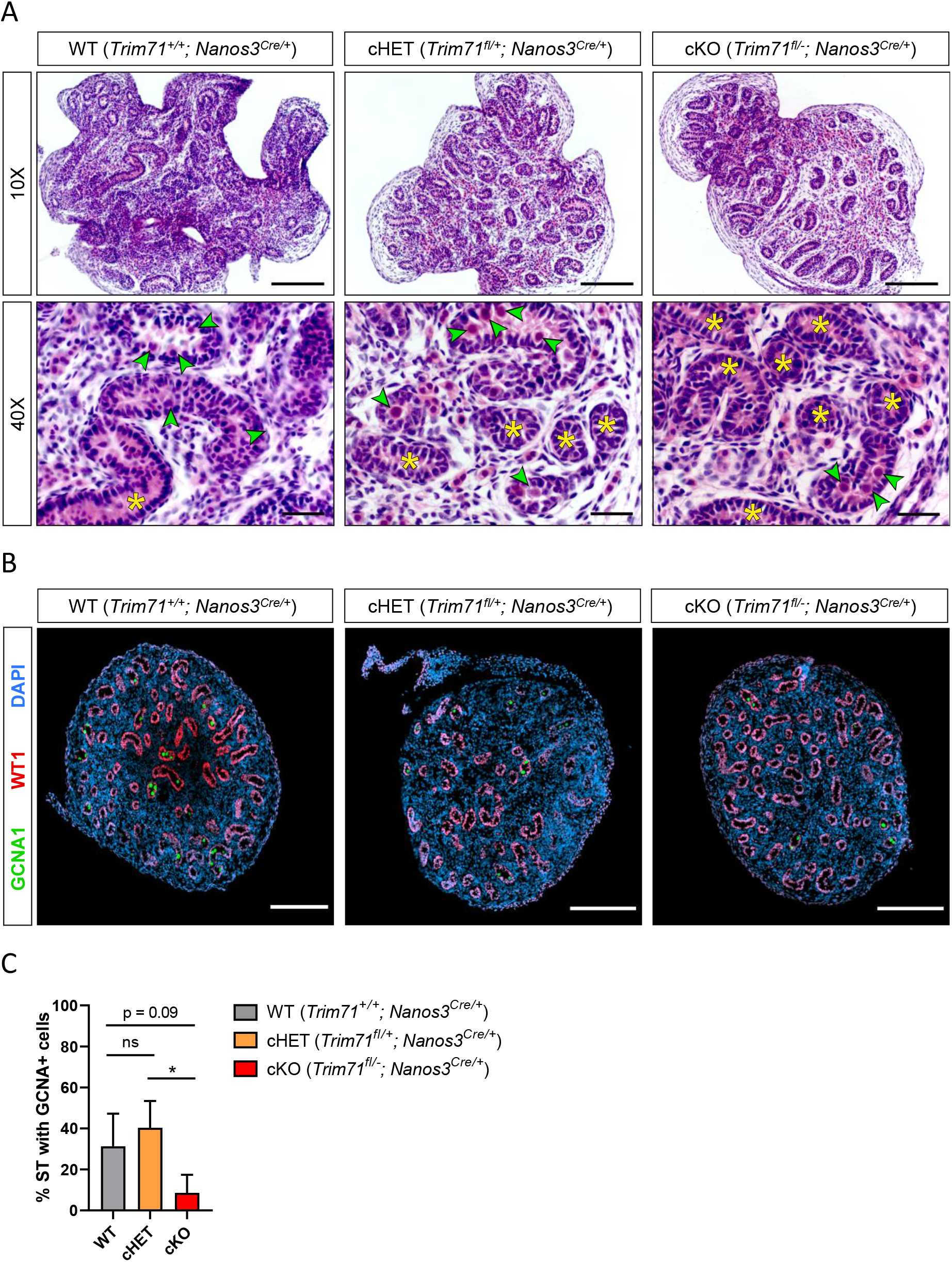
Infertility in germline-specific *Trim71* cKO male mice has an embryonic origin. **A)** Representative H&E stainings on paraffin testes cross-sections from neonatal (P0.5) wild type (WT, *Trim71*^+/+^; *Nanos3*^Cre/+^), germline-specific *Trim71* heterozygous (cHET, *Trim71*^fl/+^; *Nanos3*^Cre/+^) and germline-specific *Trim71* knockout (cKO, *Trim71*^fl/-^; *Nanos3*^Cre/+^) male mice. Gonocytes within the seminiferous tubules are marked with a green arrow head, and gonocyte-lacking seminiferous tubules are marked with a yellow asterisk (*) in 40x magnification images. Scale bars represent 100 μm and 20 μm in in 10x and 40x magnifications, respectively. **B)** Representative immunofluorescence stainings on testes cryo-sections from neonatal (P0.5) wild type (WT, *Trim71*^+/+^; *Nanos3*^Cre/+^), germline-specific *Trim71* heterozygous (cHET, *Trim71*^fl/+^; *Nanos3*^Cre/+^) and germline-specific *Trim71* knockout (cKO, *Trim71*^fl/-^; *Nanos3*^Cre/+^) male mice. Images show co-staining with GCNA1 (gonocytes), WT1 (Sertoli cells) and DAPI (nuclei). Scale bars represent 200 μm. **C)** Quantification of seminiferous tubules (ST) containing GCNA+ cells per testis cross-section from B, depicted as percentages. Error bars represent SD (n=3). *P-value < 0.05, ns = non-significant (unpaired Student’s t-test). See also Suppl. Fig. 2.

### TRIM71 is required for the optimal differentiation of ESCs into PGCs *in vitro*

In order to determine whether *Trim71* deficiency causes PGC specification defects, wild type (WT, *Trim71*^*fl/fl*^) and *Trim71* knockout (KO, *Trim71*^*-/-*^) murine ESCs^28^ were differentiated into PGC-like cells (PGCLCs) *in vitro* as previously described^46^. In short, ESCs (d0) growing under naїve conditions were primed to EpiSCs (d2) for two days in monolayer culture and differentiated into PGCLCs (d8) for six more days in the presence of BMP4 while growing as spheroids (Fig. 4A). ESCs (d0) represent *in vivo* embryonic stem cells of the blastocyst’s inner cell mass at E3.5-4.5 while EpiSCs (d2) represent *in vivo* epiblast stem cells at E5.5-6.5, and PGCLCs (d8) should correspond to *in vivo* early PGCs at E7.5^46^. Thus, several markers were measured in bulk cell populations at d0, d2 and d8 to monitor the differentiation process (Suppl. Fig. 3).

**Figure 4.**
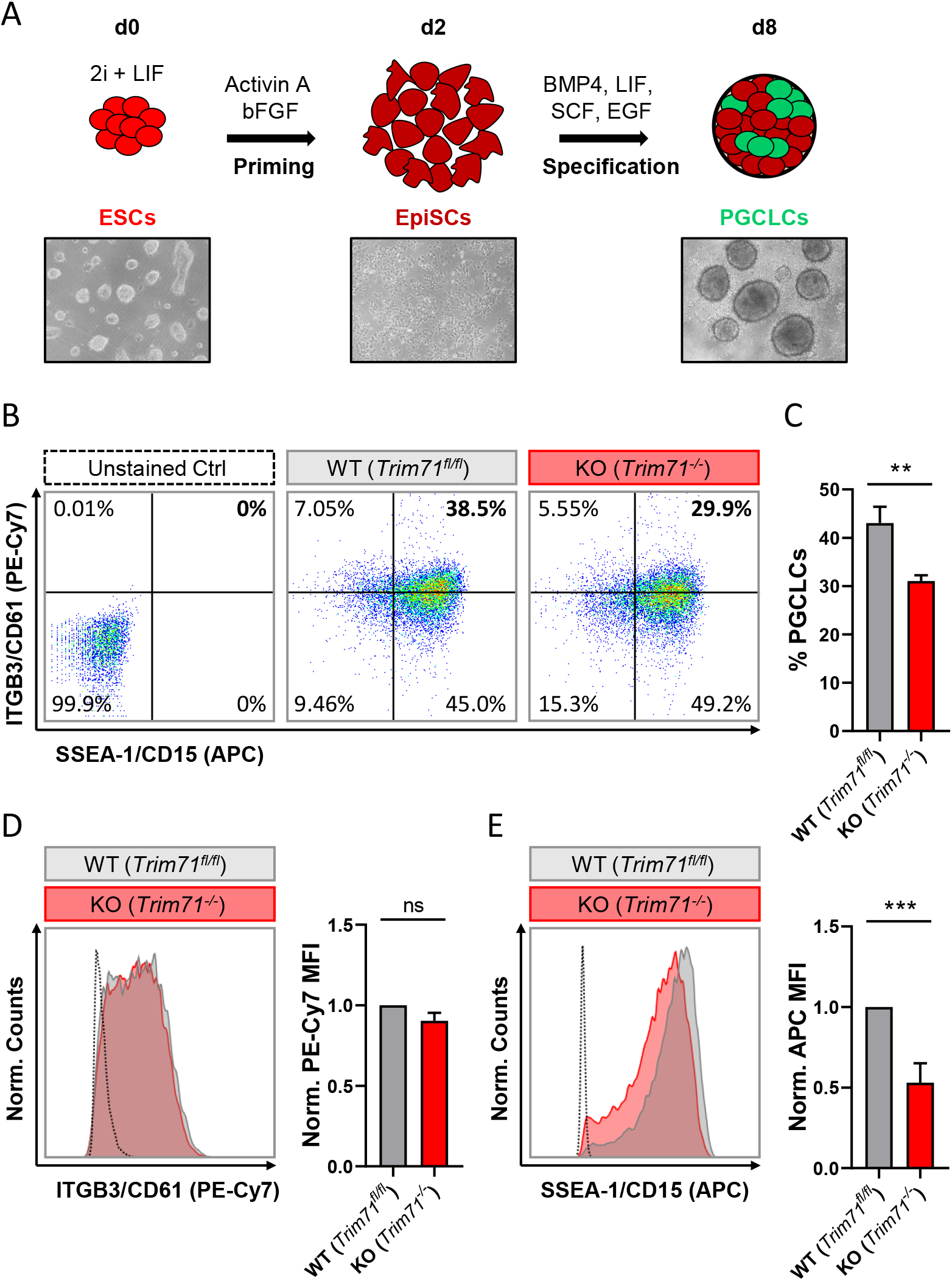
*Trim71*-deficient mouse ESCs show an impairment during PGC specification. **A)** Schematic representation of the methodology for the *in vitro* differentiation of murine ESCs into PGC-like cells (PGCLCs) accompanied by representative microscopic images (10x magnifications) of cells at different stages of the differentiation process (d0, d2 and d8). **B)** Representative flow cytometry scatter plots from d8 cells derived from wild type (WT, *Trim71*^*fl/fl*^) and *Trim71* knockout (KO, *Trim71*^*-/*-^) mouse ESCs stained for the PGC-specific surface markers ITGB3/CD61 (PE-Cy7) and SSEA-1/CD15 (APC). Double positively stained cells are considered PGCLCs. **C)** Quantification of PGCLCs in several experimental replicates of B. Error bars represent SEM (n=6). **D)** Representative histograms for individual ITGB3/CD61 (PE-Cy7) and **E)** SSEA-1/CD15 (APC) staining of d8 cells derived from wild type (WT, *Trim71*^*fl/fl*^) and *Trim71* knockout (KO, *Trim71*^*-/*-^) murine ESCs accompanied by the median fluorescence intensity (MFI) quantification for the respective staining in replicate experiments. Error bars represent SEM (n=6). ***P-value < 0.005; **P-value < 0.001; ns = non-significant (unpaired Student’s t-test). See also Suppl. Fig. 3.

Both wild type and *Trim71*-deficient ESCs were primed to EpiSCs as shown by a decrease of the naїve pluripotency marker *Klf4*^46,47^ and a simultaneous increase of the primed pluripotency epiblast-specific marker *Dnmt3b*^46,48^ at d2 (Suppl. Fig. 3A-B). The subsequent *Dnmt3b* downregulation observed at d8 was indicative of a successful specification induction by BMP4, as the early PGC marker *Prdm14* represses *Dnmt3b* to enable PGCs epigenetic reprogramming^49,50^. Accordingly, both wild type and *Trim71*-deficient cells showed a significant upregulation of PGC-specific markers at d8 compared to d2 including *Prdm14, Prdm1/Blimp1* and their downstream targets *Tfap2c* and *Nanos3* (Suppl. Fig. 3C-F). However, the levels of *Prdm1/Blimp1* at d8 were significantly lower in *Trim71*-deficient cells than in wild type cells, an effect that was also apparent – although not significant – for *Nanos3*, suggesting a moderate impairment in the specification of *Trim71*-deficient PGCLCs. Importantly, while an upregulation of the early PGC marker *Nanos3*^11^ was already observable at d8, the expression of the late PGC marker *Ddx4/Vasa*^14,15^ was not increased (Suppl. Fig. 3G), indicating that the *in vitro*-generated PGCLCs represent *in vivo* pre-migratory PGCs at the end of the specification phase (E7.5), as expected.

In order to estimate the number of PGCLCs specified from wild type and *Trim71*-deficient ESCs, d8-bulk populations were stained for the surface markers ITGB3/CD61 (PE-Cy7) and SSEA-1/CD15 (APC) – previously reported to unequivocally identify PGCLCs^46^ – and were analyzed by flow cytometry. We found that the number of PGCLCs derived from *Trim71*-deficient ESCs was reduced by about 25% (Fig. 4B-C). A significant decrease of the PGC-specific marker SSEA-1/CD15 (APC) – but not of ITGB3/CD61 (PE-Cy7) – was also observed in PGCLCs derived from *Trim71*-deficient ESCs (Fig. 4D-E).

Of note, *Oct4* is required for the maintenance of PGCs after their specification^51^. The highest expression of *Oct4* in the mouse embryo is observed in the early blastocyst and decreases progressively in the epiblast upon gastrulation until being restricted to PGCs at E7.5^20,52^. Accordingly, we detected a high *Oct4* expression in ESCs (d0) which significantly decreased upon priming to EpiSCs (d2) and was maintained in PGCLCs (d8) upon specification (Suppl. Fig. 3H). A similar pattern was observed for *Trim71* expression in wild type cells (Suppl. Fig. 3I) with a decrease in its mRNA level after priming (d2) and a sustained expression in PGCLCs (d8). While our results have revealed a moderate impairment in the specification of *Trim71*-deficient PGCLCs, we believe that this effect may not fully explain the dramatic phenotype observed in *Trim71* cKO mice. This, together with the sustained *Trim71* expression in PGCLCs at d8 suggest a further role for *Trim71* downstream of PGC specification.

### TRIM71 controls proliferation in the germline-derived tumor cell line NCCIT

After specification (E7.5), PGCs migrate and colonize the genital ridges. PGCs then actively proliferate before undergoing sex determination (E10.5), followed by mitotic arrest (E13.5) in case of male PGCs – now termed gonocytes. Thus, the cluster of ~40 founder PGCs grows to approximately 25,000 gonocytes in the developing male gonads during this period^12,13,17,18^. Importantly, the failure of male PGCs to further differentiate into gametes can lead to the development of testicular GCT (TGCT)^2–4^.

TRIM71 is highly expressed in several GCT-derived cell lines^25,29,33^ (Suppl. Fig. 4A) and found to be upregulated in TGCT patients^53^ (Supp. Fig. 4B). We therefore used the GCT-derived embryonic carcinoma cell line NCCIT to generate *TRIM71* frameshift mutations via CRISPR/Cas9 and evaluated whether TRIM71 controls NCCIT proliferation in growth competition assays. *TRIM71* mutations were generated by using two different single guide RNAs (sgRNA), one targeting the N-terminal RING domain and another targeting the C-terminal NHL domain (Suppl. Fig. 4C). For the RING sgRNA (△RING), the generation of single NCCIT clones showed that an 83 kDa N-truncated RINGless version of TRIM71 is generated from the use of an alternative in-frame ATG codon present downstream of the targeting region (Suppl. Fig. 4D). For the NHL sgRNA (△NHL6), generation of single clones showed that the resultant protein was either absent or had a C-terminal truncation of the last NHL repeat (Suppl. Fig. 4E), a mutation which is already known to mimic the full KO phenotype *in vivo*^31^ and *in vitro*^29^.

Single cell clones often have a selection bias towards robust *in vitro* proliferation and survival. In order to study TRIM71-dependent cell proliferation in an unbiased manner, we analyzed cell population dynamics during *in vitro* growth competition assays using non-clonal mixed pools of wild type and *TRIM71* mutant cells. To this end, NCCIT cells were mock-transfected with an empty vector (EV) or with TRIM71-targeting vectors (△RING or △NHL6) and sorted for Cas9-GFP+ cells. Bulk-transfected EV (wild type) and △RING/△NHL6 (*TRIM71* mutant) NCCIT pure populations (Suppl. Fig. 5A-B) were mixed in a 1:1 ratio. We then evaluated changes in the distribution of single reads via NGS (Illumina MiSeq system)^54^ for wild type and *TRIM71* mutant alleles over a time period of 21 days and classified them as wild type reads, reads with *TRIM71* frameshift (loss-of-function) mutations and reads with *TRIM71* in-frame mutations (Fig. 5). As a negative control, pure wild type NCCIT (EV) cell populations were analyzed at day 0 and day 21, showing no relevant changes in allele distribution (Suppl. Fig. 5C-D). In contrast, the percentages of alleles with *TRIM71* frameshift mutations dropped ∼2-fold (from 11.34 % to 5.96 %) for NCCIT ΔRING cells within 21 days (Fig. 5A) and ~13-fold (from 32.46 % to 2.53 %) for NCCIT ΔNHL6 cells in the same time period (Fig. 5B). Interestingly, *TRIM71* in-frame mutations also resulted in a growth disadvantage, although to a lesser degree in each case (Fig. 5A-B).

**Figure 5.**
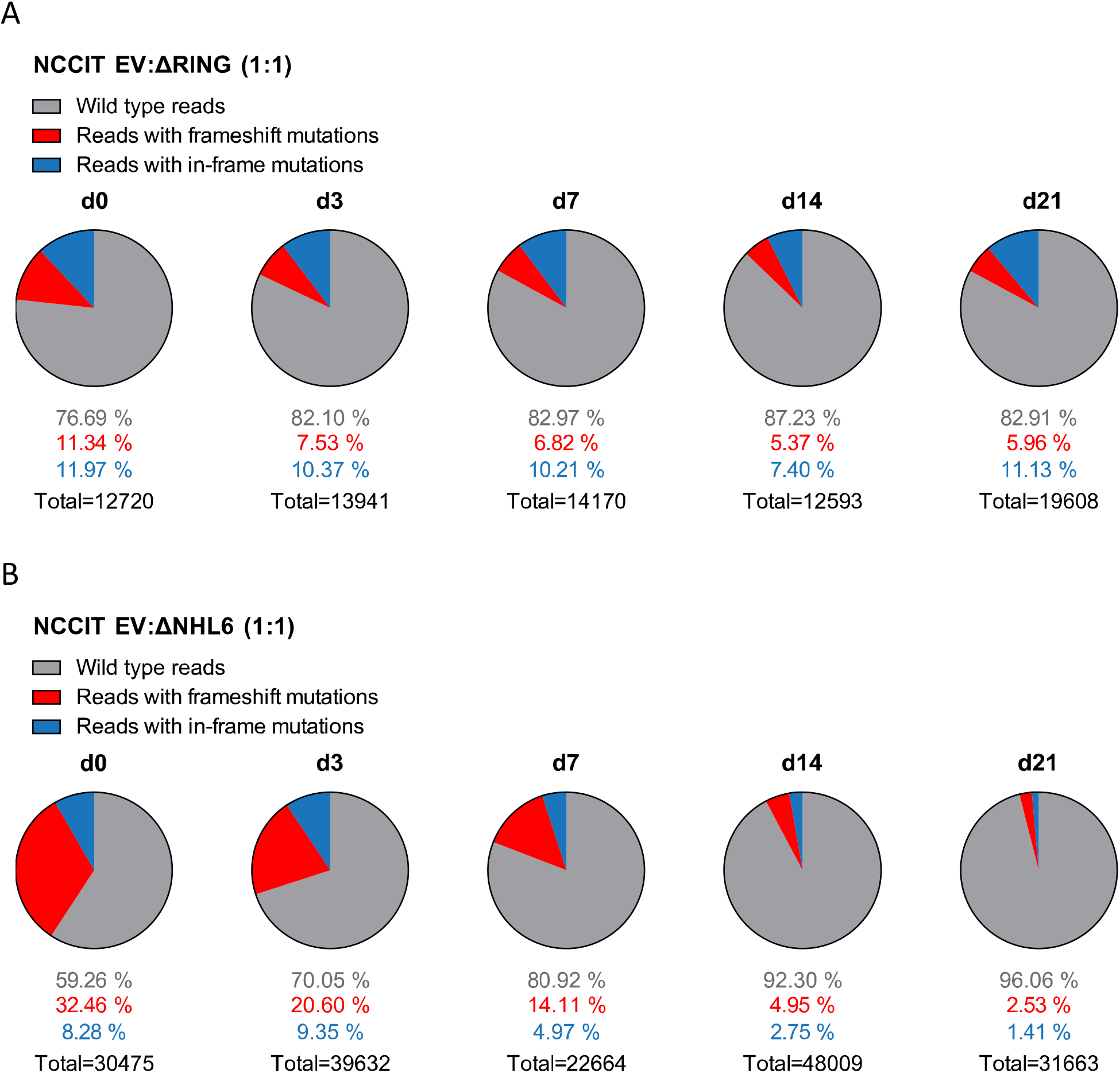
NCCIT cells with *TRIM71* mutations show proliferation defects in growth competition assays. **A)** Pie charts showing allele frequencies for wild type (EV) and TRIM71 RING mutant (ΔRING) or **B)** TRIM71 NHL mutant (ΔNHL6) NCCIT cells obtained at different time points during growth competition assays. For the assay, NCCIT wild type cells (EV) were mixed 1:1 with either TRIM71 RING mutant (ΔRING) or TRIM71 NHL mutant (ΔNHL6) NCCIT cells and directly analyzed (d0) before their culturing, following by subsequent analysis at several time points (d3, d7, d14 and d21). Allele frequency was analyzed via NGS using the Illumina MiSeq platform. For *TRIM71* mutant alleles, reads with in-frame mutations and frameshift mutations (loss-of-function) in each respective domain (RING or NHL) are depicted. The total number of sequencing reads for each time point is indicated under each respective pie chart. See also Suppl. Fig. 4 and 5.

These results demonstrate that TRIM71 is involved in the control of proliferation in NCCIT cells, consistent with previous studies involving other tumor cell lines^22,24,29,55^. Our work, together with the elevated *TRIM71* expression observed in TGCT patients, confirms a role for TRIM71 supporting the proliferation/maintenance of TGCT. Considering that PGCs undergo active proliferation after reaching the genital ridges, and that TGCT derive from PGCs which fail to further develop into gonocytes, the deficit of germ cells observed in our *Trim71* cKO male mouse possibly results from a combination of specification defects and subsequent proliferation defects in PGCs upon TRIM71 ablation.

### Exome sequencing data identifies *TRIM71* variants in infertile men with severely impaired spermatogenesis

The causes of the SCO phenotype in men remain poorly understood^43–45^. In order to identify novel associated genes, we utilized exome sequencing data of 247 SCO subjects belonging to the Male Reproductive Genomics (MERGE) study. We employed our in-house software Sciobase and our newly developed software Haystack to analyze loss-of-function (LoF) variants. After strict filtering based on quality criteria, minor allele frequency (MAF) in the general population, high expression in human testes (GTEx) and absence of LoF variants in individuals with complete spermatogenesis, we identified a total of 721 genes with LoF variants specifically found in SCO patients (Fig. 6A-C). Via this independent and unbiased approach, we found *TRIM71* to be present among these genes, revealing a possible association with human male infertility. This finding is supported by our previous data showing that TRIM71 deficiency results in an SCO-like phenotype in male mice. Of note, most of the identified human genes carried LoF variants in only one patient within the studied cohort and none of the identified genes carried LoF variants in more than four individuals, indicating that the SCO phenotype is highly heterogeneous (Fig. 6B). Similar results were observed upon inclusion of genes with lower expression in the testis (Fig. 6C).

**Figure 6.**
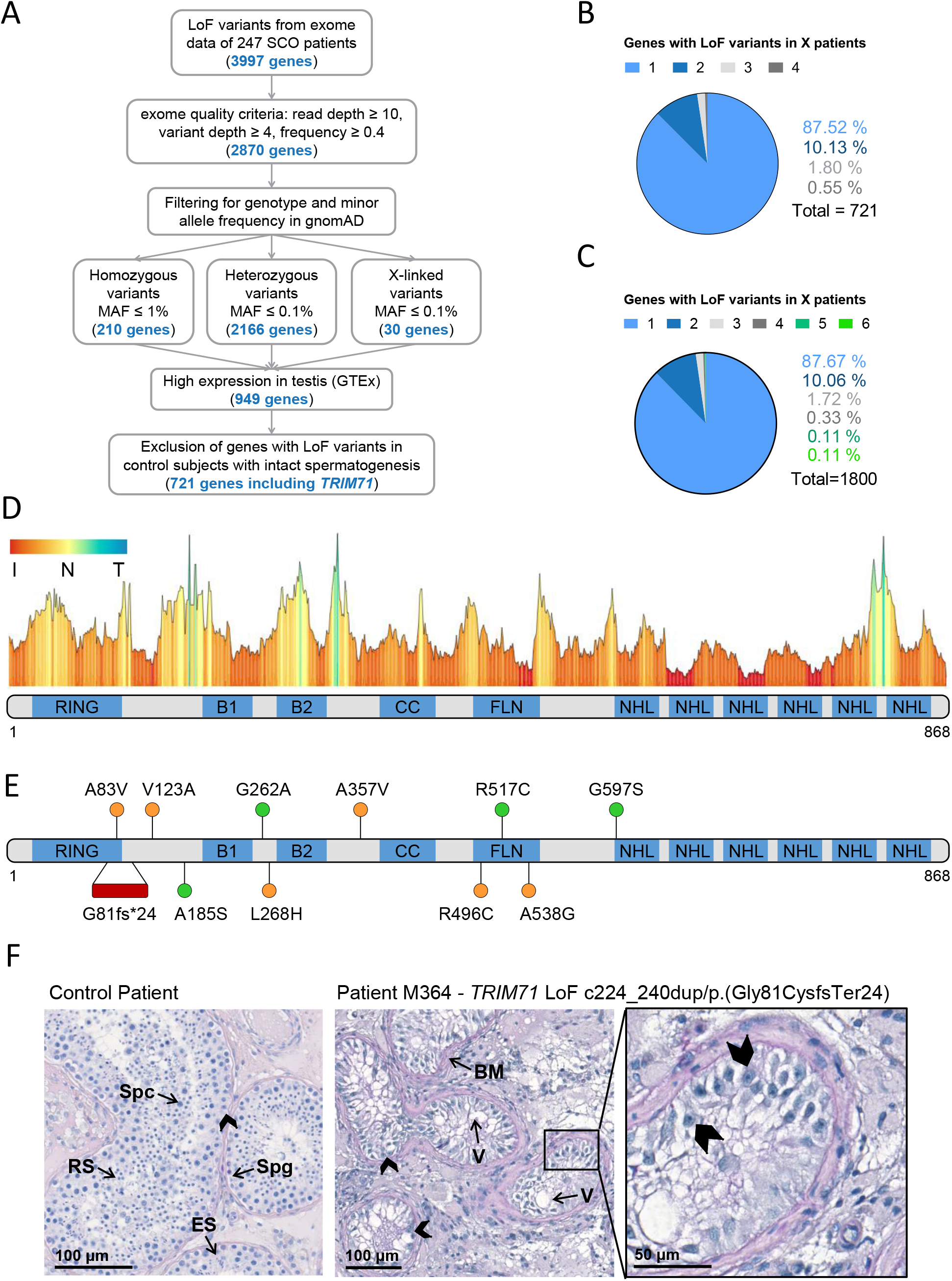
Exome sequencing in infertile SCO patients reveals an association of *TRIM71* with human male infertility. **A)** Filtering scheme detailing gene prioritization of exome sequencing data from 247 individuals with Sertoli cell-only (SCO) phenotype using Sciobase and Haystack (see details in the Methods section). Gene numbers remaining after each filtering step are indicated in blue. Loss-of-function (LoF) variants include stop, frameshift, splice acceptor and splice donor variants. gnomAD = Genome Aggregation Database; MAF = minor allele frequency; GTEx = Genotype-Tissue Expression Portal. **B)** Percentages of genes (total = 721, obtained at the end of the filtering process depicted in A) affected by LoF variants in 1-4 SCO patients. None of the genes carried LoF variants in more than 4 patients in the SCO cohort (n = 247). **C)** Percentages of genes (total = 1800, obtained at the end of the filtering process depicted in A, excluding “High expression in testis - GTEx” filtering step) affected by LoF variants in 1-6 SCO patients. None of the genes carried LoF variants in more than 6 patients in the SCO cohort (n = 247). **D)** TRIM71 mutation tolerance landscape obtained from MetaDome (https://stuart.radboudumc.nl/metadome/). The graph displays missense over synonymous variant ratios per position for the entire protein, based on gnomAD variants. I = intolerant; N = neutral; T = tolerant. **E)** Schematic representation of TRIM71 domain structural organization depicting the location of the genetic variants identified in MERGE. Red = presumably pathogenic LoF variants; orange = missense variants of uncertain significance; green = missense variants found also in proven fathers. **F)** Periodic acid–Schiff stainings of testis sections obtained by testicular biopsy from a control patient and an SCO patient (subject M364), who harbors the *TRIM71* variant c.224_240dup/p.(Gly81CysfsTer24) in heterozygosis. The control section shows intact spermatogenesis (Spg = spermatogonia; Spc = spermatocytes; RS = round spermatids; ES = elongated spermatids) as opposed to subject M364, whose histological evaluation revealed an SCO phenotype (exemplary Sertoli cells indicated by arrow heads) with several degenerated seminiferous tubules, frequently including large vacuoles (V) and thickened basement membranes (BM). See also Suppl. Fig. 6.

For a further evaluation of *TRIM71* as a candidate gene associated with human male infertility, we screened 908 infertile patients - including those with SCO from MERGE - for variants in *TRIM71*. We identified 11 different rare (MAF ≤ 0.001) heterozygous *TRIM71* variants, including the aforementioned LoF variant and 10 novel missense variants, present in a total of 14 infertile individuals with severely impaired spermatogenesis and varying histological phenotypes (Fig. 6D-F and Table 1). The majority of subjects (11 out of 14) were azoospermic, and some of them presented with reduced testicular volumes, elevated luteinizing hormone (LH) levels, elevated follicle-stimulating hormone (FSH) levels and/or reduced serum testosterone (T) levels (Table 1), all signs of broad testicular dysfunction present in 60% of azoospermic men^44^.

**Table 1:**
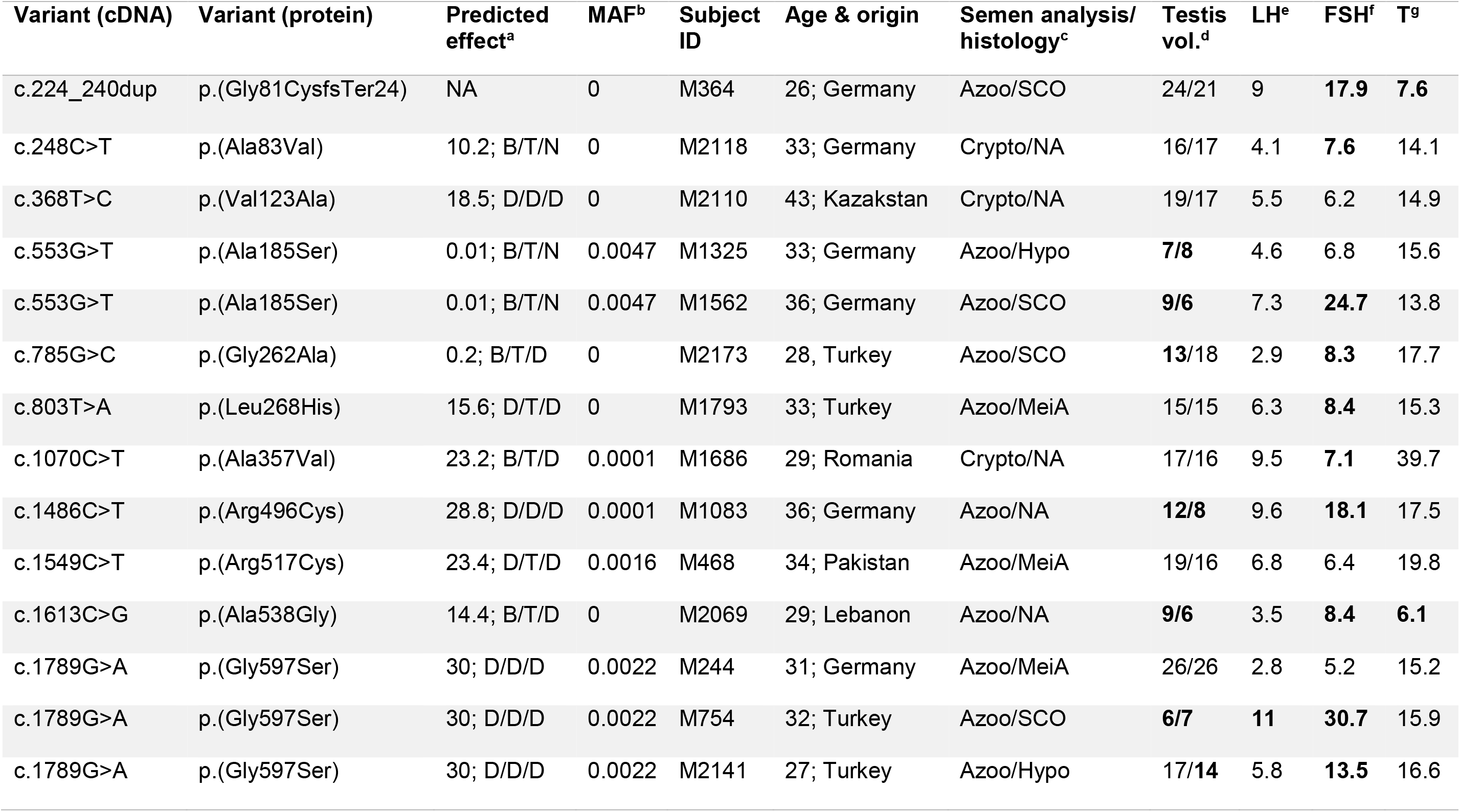
*TRIM71* variants identified in infertile men within the MERGE cohort. ^a^Predicted effect of each *TRIM71* variant estimated by four different pathogenicity prediction algorithms (CADD/PolyPhen2/SIFT/MutationTaster, shown in the same order). For CADD, variants with values above 20 are more likely to be deleterious to protein function. For the rest, D = damaging/deleterious; T = tolerated; B = benign; N = neutral; NA = not available. ^b^MAF (minor allele frequency) values derive from gnomAD and are considered rare if MAF < 0.001 when occurring in heterozygosis. ^c^Semen analysis was performed for all patients (Azoo = azoospermia; Crypto = cryptozoospermia), and if biopsies were available, patients’ phenotypes were also histologically assessed (SCO = Sertoli cell-only syndrome; MeiA = meiotic arrest; Hypo = hypospermatogenesis). ^d^Testicular volumes (testis vol., right/left, ref. >15 mL each; bold = values outside the normal range). ^e^Luteinizing hormone (LH, ref. 2-10 IU/L; bold = values outside the normal range). ^f^Follicle-stimulating hormone (FSH, ref. 1-7 IU/L; bold = values outside the normal range). ^g^Testosterone (T, ref. >12 nmol/L; bold = values outside the normal range).

Generally, *TRIM71* seems to be rather intolerant to missense variation (Figure 6D)^56^. This was also indicated by a high Z score value of 3.28 for *TRIM71*. The Z score is a metric computed by gnomAD database and ranges from -5 to 5, with higher values indicating an intolerance to variation and, therefore, a higher likelihood for *TRIM71* variants to disrupt TRIM71 function. Furthermore, parameters such as MAF or pathogenicity prediction algorithms were used to estimate the reliability for the association of each variant with male infertility (Table 1). The conservation of the affected residues for each variant was also evaluated (Suppl. Fig. 6A).

Four of the *TRIM71* missense variants identified in infertile men were also found in control subjects with complete spermatogenesis (n = 89) or in a Dutch cohort of proven fathers (n = 5784), making their association with male infertility rather unlikely (Fig 6E, green). In contrast, other identified *TRIM71* missense variants were present in one patient each but neither in control subjects nor in the Dutch cohort of proven fathers (Fig 6E, orange). Of those, c.368T>C/p.(Val123Ala), c.803T>A/p.(Leu268His), c.1070C>T/p.(Ala357Val) and c.1486C>T/p.(Arg496Cys) were considered damaging by at least two out of four pathogenicity prediction algorithms (Table 1), and the respective residues for those variants were highly conserved among vertebrates (Suppl. Fig 6A). Thus, these variants have a higher likelihood of being associated with male infertility. As an additional line of evidence, all patients carrying variants in *TRIM71* (Table 1) were evaluated for relevant variants in other genes previously reported in association with male infertility (n=181, listed in Suppl. Table 1). Importantly, one heterozygous frameshift variant was found in *SYCP2* in patient M1686, who carries the *TRIM71* variant c.1070C>T/p.(Ala357Val). *SYCP2* was recently associated with male infertility^57^, and it is thus unlikely that this patient’s *TRIM71* variant alone is responsible for his cryptozoospermia, but an oligogenic cause of his phenotype could be hypothesized. Furthermore, a heterozygous missense variant for the gene *NNT* was identified in patient M1083, who carries the *TRIM71* variant c.1486C>T/p.(Arg496Cys). In this case, the *NNT* missense variant is of unclear significance according to ACMG/AMP guidelines, and its functional contribution to the patients’ phenotype can therefore neither be confirmed nor excluded at this point. Collectively, our analysis leaves the *TRIM71* variants c.368T>C/p.(Val123Ala) and c.803T>A/p.(Leu268His) as the likeliest candidates for being associated with male infertility. Nevertheless, the functional effects of all *TRIM71* missense variants should be evaluated experimentally in the future in order to definitively determine their pathogenicity.

### Exome sequencing data reveals an association of *TRIM71* LoF variants with the SCO phenotype

As mentioned above, a *TRIM71* LoF variant - c.224_240dup/p.(Gly81CysfsTer24) - was identified via exome sequencing in subject M364, who presented with a clear SCO phenotype (Fig. 6E-F and Table 1). This variant consists of a duplication of 17 nucleotides which generates a frameshift resulting in a premature translation termination downstream of the RING domain (Suppl. Fig. 6B). This particular *TRIM71* LoF variant has not been described in any public database so far, and was classified as likely pathogenic according to ACMG/AMP guidelines. Furthermore, the observed/expected (o/e) score computed for *TRIM71* in the gnomAD database was extremely low (o/e score = 0.04, 90% confidence interval 0.01-0.17). The o/e score compares the number of observed versus theoretically expected heterozygous LoF variants for a gene of interest. Without any selection pressure applied, an o/e ratio of around 1 would be expected for any given gene, whereas values below 0.35 are a rather clear sign of selection pressure against LoF variants, likely leading to haploinsufficiency intolerance.

Furthermore, subject M364 carries neither additional LoF variants nor likely pathogenic missense variants in any other known infertility-associated genes (listed in Suppl. Table 1). This patient presented with normal testicular volumes and LH levels, but increased FSH and decreased testosterone levels (Table 1), as is often the case in men with impaired spermatogenesis. Of note, no *TRIM71* LoF variants were found in the Dutch cohort of proven fathers, and only one LoF variant was present in the exome data from 125,748 presumably healthy individuals from gnomAD^58^. This constitutes a significant enrichment of *TRIM71* LoF variants in the MERGE cohort (n = 908; p = 0.01) and an even higher enrichment in the SCO subcohort (n = 247; p = 0.004), supporting a reliable association of *TRIM71* LoF variants with human male infertility.

## Discussion

TRIM71 is a stem cell-specific protein expressed early in development and with an essential function for embryogenesis^30–33^. Embryonic lethality in *Trim71* full knockout mice is accompanied by neural tube closure defects, revealing also a function of TRIM71 in the development of the nervous system^30–33^. Indeed, TRIM71 is known to promote self-renewal of neural progenitor cells^21^, and *TRIM71* variants have been associated with human congenital hydrocephalus, a brain developmental disease characterized by enlarged brain ventricles due to an abnormal accumulation of cerebrospinal fluid^59^. Furthermore, TRIM71 has been connected with carcinogenesis in several studies^22,24,29,55,60^. Altogether, previous research on TRIM71 has been mostly focused on embryonic and neural stem cells as well as cancer cells. Our work here reports a novel role for TRIM71 in the generation and maintenance of germ cells during embryonic gonad development with crucial implications in murine and human fertility.

In line with previous studies^33,34^, our work reveals an expression of *Trim71* in adult mouse testes which we found to be confined to SSCs (Fig. 1). We showed that germline-specific ablation of *Trim71* early in mouse development causes a substantial reduction of gonad size and infertility in both sexes (Fig. 2). Further characterization of adult *Trim71* cKO mouse testes revealed a significant downregulation of SSC-specific and spermatid markers caused by a strong reduction in the number of developing germ cells (Fig. 2), a condition which in humans is known as Sertoli cell-only (SCO) phenotype^43–45^. Via exome sequencing of human infertile patients, we uncovered an association of *TRIM71* with the SCO phenotype (Fig. 6).

In humans, infertility affects 10-15 % of couples trying to conceive^4^. Male factors contribute in about half of couples and usually genetic causes correlate with a more severe spermatogenic impairment^61^. Lately, several genes have been identified as monogenic causes for azoospermia due to meiotic arrest (e.g. TEX11^62^, STAG3^63^, M1AP^64^ and SHOC1^65^). However, the SCO phenotype seems to be a highly heterogeneous condition, as indicated by our data, making the identification of monogenic causes specially challenging. Our work uncovers *TRIM71* as the first monogenic cause for this condition, based on the SCO-like phenotype observed in our germline-specific *Trim71* cKO mouse together with the *TRIM71* LoF variant that we found in a SCO patient. The finding of *TRIM71* as a novel SCO candidate gene is relevant in terms of patient care, as it may allow future patients to be provided with a causal diagnosis for their azoospermia, and the potential success of testicular biopsy and sperm extraction with the aim to perform *in vitro* fertilization could be predicted in advance.

In contrast to the convincing relevance of the *TRIM71* LoF variant, assessing the role of the identified rare *TRIM71* missense variants in infertile patients of varying histological phenotypes is much more challenging. Of note, variants in the same gene may cause a spectrum of histological phenotypes^64^. Although our analysis provides a prediction for the pathogenicity of these variants, their functional effects on human male fertility are yet of unclear significance and can only be determined experimentally. Nevertheless, it is worth emphasizing that most missense variants identified in *TRIM71* cluster outside of the NHL domain, as mutations affecting the NHL domain have been proven to be highly deleterious^66,67^ and compromise survival during early embryogenesis^31^. Future experiments using these *TRIM71* variants should determine whether they functionally affect TRIM71’s E3 ubiquitin ligase role, and if so, how exactly TRIM71-mediated ubiquitylation contributes to germ cell development.

Our germline-specific *Trim71* cKO mouse model provided further evidence for the role of *Trim71* in germ cell development. The SCO-like phenotype was already apparent in the testes of neonatal (P0.5) cKO mice, indicating that TRIM71-induced germline defects have an embryonic origin (Fig. 3). However, early developmental defects are likely amplified during postnatal mitotic reactivation and pubertal spermatogenesis as described by a recent work showing infertility in a different germline-specific *Trim71* knockout (*Trim71*^*-/fl*^; *Ddx4*^*Cre/+*^) mouse model^34^. In contrast to our observations in *Trim71*^*fl/-*^; *Nanos3*^*Cre/+*^ mice, defects in *Trim71*^*-/fl*^; *Ddx4*^*Cre/+*^ mice were only detected in pubertal (P10) and adult (P56) mice, with germ cell numbers not yet altered in neonatal (P1) mice^34^. This discrepancy may result from an earlier Cre recombinase expression in our cKO model, since *Nanos3* is expressed during PGC specification (E7.5)^5,10,11^, and *Ddx4* is expressed after colonization of the genital ridges (E10.5)^14,15^. In fact, *Ddx4*-induced recombination was previously reported to occur even after sex determination as late as E15.0^68^. This might also explain why *Trim71*^*fl/-*^; *Ddx4*^*Cre/+*^ females are fertile (Prof. Xin Wu, personal communication, June 22, 2020), while we found *Trim71*^*fl/-*^; *Nanos3*^*Cre/+*^ females to be sterile. Notably, studies in *C. elegans* showed that the nematode homolog of TRIM71, LIN-41, is required for normal oocyte growth and meiotic maturation, with *lin-41* depletion causing sterility in females^69,70^. Further studies are required to characterize the function of TRIM71 in mammalian female gonad development and fertility.

Although we do not exclude sex-specific functions of TRIM71 in the germline, we found both male and female *Trim71* cKO mice to be infertile, indicating that germ cell developmental defects may occur prior to sex determination of PGCs. Indeed, we observed PGC specification defects upon *in-vitro* differentiation of *Trim71*-deficient ESCs into PGCLCs, with reduced numbers of PGCLCs at the end of the specification process and decreased expression of the PGC-specific markers *Prdm1/Blimp1, Nanos3* and SSEA-1/CD15 upon TRIM71 depletion (Fig. 4). Interestingly, *Trim71* expression was still remarkably high in PGCLCs, suggesting its further involvement in subsequent stages of germ cell development. In fact, PGC specification defects may be followed by defects in the maintenance of the PGC pool during the stages of migration and genital ridge colonization, as we found that TRIM71 controls proliferation/self-renewal in the germline-derived tumor cell line NCCIT, which, to some extent, represent proliferative PGCs^71^.

The capability of self-renewal in a cell population reflects the balance between cell proliferation, differentiation and apoptosis. After PGC colonize the genital ridges, a significant expansion of the PGC pool occurs^12,13,17,18^. Furthermore, mitotically arrested male PGC undergo several massive apoptotic waves within the developing gonads: in mice, the first one occurs between E13.5 and E17.0 and is followed by a second one around birth and a third during the first wave of spermatogenesis between P10 and P13^72^. Therefore, the role of TRIM71 in the maintenance of the PGC pool during these phases may extend beyond proliferation control, as previous studies have shown the involvement of TRIM71 not only in the control of cell proliferation, but also during cell differentiation and apoptosis. For instance, TRIM71 represses the mRNA of several cell cycle inhibitors, including *CDKN1A, E2F7* and *Rbp1/2*, promoting proliferation of ESC and several cancer cell lines^25,26,29^. Proliferation of ESCs, neural progenitor cells, lung cancer cells and liver cancer cells was also promoted via TRIM71’s E3 ubiquitin ligase function^21,22,55,73^. TRIM71 has also been shown to prevent apoptosis in developmental and oncogenic processes, specifically via p53 ubiquitylation and proteasomal degradation in breast cancer cells^24^ and ovarian cancer cells^60^ as well as in differentiating ESCs during mouse brain development^23^. More importantly, a recent study described an impaired proliferation and an increased apoptosis in primary murine SSCs upon TRIM71 knockdown^34^. Collectively, these findings together with our growth competition assays in NCCIT cells, support a role for TRIM71 in the control of proliferation and possibly also apoptosis.

Finally, premature differentiation of TRIM71-deficient cells could be also contributing to the germ cell loss observed in our *Trim71* cKO mice. A previous study has shown that *Trim71*-deficient ESCs are primed towards neural differentiation^28^. TRIM71 is also known to repress several mRNAs promoting differentiation, such as *HOXA5*^26^ or *EGR1*^27^. Furthermore, *Trim71*-deficient ESCs showed a premature upregulation of differentiation-promoting micro-RNAs (miRNAs), including brain-specific and gonad-specific miRNAs^28^, highlighting a yet-unknown role for TRIM71-mediated miRNA regulation in early gonad development. Which mRNA, miRNA and/or protein targets are precisely regulated by TRIM71 to fulfill its functions in the germline remains to be further investigated.

In summary, we have shown that germline-specific *Trim71* cKO male mice are infertile with testes predominantly displaying an SCO-like phenotype. We have identified *TRIM71* variants in infertile men with severely impaired spermatogenesis, including a LoF variant in an SCO patient. The SCO-like phenotype was already apparent in neonatal P0.5 *Trim71* cKO mice. Our *in vitro* assays suggested that germ cell deficiency in these mice may result from combined defects in the specification and maintenance of the PGC pool during embryonic gonad development. In a similar fashion, TRIM71 may support the maintenance of embryonic carcinoma cells and promote the development of GCT. Future unravelling of the molecular mechanisms by which TRIM71 governs germ cell biology will shed further light on the underlying causes of infertility. A deeper understanding of these processes will contribute to the development of new diagnostic and therapeutic strategies for reproductive medicine and for the treatment of GCT.

## Methods

### Mouse generation

All animal experiments were conducted in a licensed animal facility in accordance with the German law on the protection of experimental animals (the German animal welfare act), and were approved (approval number 87-51.04.2011.A063) by local authorities of the state of Nordrhein-Westfalen (Landesamt für Natur-, Umwelt-und Verbraucherschutz NRW).

The generation of the full *Trim71* knockout mouse (*Trim71*^*fl/fl*^; *Rosa26-CreERT2*) was previously described^28^. In order to generate a germline-specific *Trim71* knockout mouse, wild type females with floxed *Trim71* alleles (WT, *Trim71*^*fl/fl*^) were bred with male mice expressing the Cre recombinase under the control of the endogenous *Nanos3* promoter in heterozygosity (*Nanos3*^*Cre/+*^)^74^. The heterozygous *Trim71*^*-/+*^; *Nanos3*^*Cre/+*^ male offspring was then crossed with wild type *Trim71*^*+/+*^ females to produce *Trim71* wild type animals with the *Nanos3*-Cre allele in heterozygosity (WT, *Trim71*^*+/+*^; *Nanos3*^*Cre/+*^), or with wild type *Trim71*^*fl/fl*^ females to produce germline-specific heterozygous (cHET, *Trim71*^*-/+*^; *Nanos3*^*Cre/+*^) and germline-specific knockout (cKO, *Trim71*^*-/fl*^; *Nanos3*^*Cre/+*^) animals.

### Genomic DNA extraction and genotyping

Genomic DNA was extracted from adult mice tail or ear biopsies or from mouse embryo yolk sacs by boiling the tissue in 50 mM NaOH for 20 minutes followed by pH neutralization of the lysate with 1/4 of 1 M Tris-Cl, pH 8.0. A three-primer strategy was used for PCR amplification of the *Trim71* locus or the *Nanos3* locus and fragments were resolved on a 2% agarose gel. Primers used are listed in Suppl. Table 2.

### Epidydmal sperm counts

Cauda epididymis were isolated from 3-month-old male mice and transferred into 0.5-1 ml of PBS pre-warmed at 35-37°C. Sperm release was achieved by performing multiple cauda incisions followed by 10-min incubation at 35-37°C to allow the sperm swim out. The sperm suspension was then diluted in water (1:15-1:40) and sperm count was determined using a Neubauer hemocytometer.

### Isolation of spermatogonial stem cells from mouse testes

Testes were isolated from 2-month-old male mice, and seminiferous tubules were exposed by removing the tunica albuginea. The tubules of several testes were pooled and incubated in approximately 10 volumes of HBSS with calcium and magnesium containing 1 mg/ml collagenase Type IV and 200 to 500 μg/ml DNAse I, followed by incubation at 37°C under gentle agitation for 15 min. The tubules were then washed three times in 10 volumes of HBSS, followed by incubation at 37°C for 5 min in HBSS containing 0.25 % trypsin and 150 μg/ml DNAse I under gentle agitation. Trypsinization was stopped by adding 20 % FBS, and the cell suspension was filtered through a 40 μm pore size nylon filter. The filtrate was centrifuged at 1000 rpm for 5 min at 4 °C and the cell pellet was resuspended in PBS before cell counting. Magnetic-activated cell sorting (MACS) was used to enrich THY1.2+ cells. To this end, 10^6^ cells were stained with 0.2 μg FITC-labelled anti-THY1.2/CD90.2 antibody (BioLegend) for 15 min at 4°C. The cells were then washed in MACS-buffer (PBS supplemented with 2 mM EDTA and 0.5% FBS) and incubated in a 1:5 dilution of anti-FITC microbeads (Miltenyi) for 15 min at 4 °C before a final washing in MACS buffer followed by centrifugation at 1000 rpm for 5 min at 4 °C. The pellet was resuspended in 500 μl MACS buffer and loaded on an AutoMACS device (Miltenyi) with the program set for positive selection. The efficiency of enrichment was 8-fold as later controlled by flow cytometry. The different cell populations were then used for RNA extraction and qRT-PCR quantification.

### H&E staining of mouse testes cross-sections

PFA-fixed paraffin-embedded murine testes cross-sections were deparaffinized by incubation at 65 °C for 15-20 min until the paraffin wax had melted, and washed in xylol twice for 10 min. Next, the sections were rehydrated by a descending ethanol dilution series (100 % (2x), 95 %, 90 %, 80 %, 70 %) for 30 sec each and were kept in distilled water until staining. Cross-sections were stained in haematoxylin for 3 min and washed in cold running tap water for 3-5 min. Afterwards, the sections were counterstained with 0.5 % eosin for 3 min, removing excess dye by rinsing in cold running tap water for 1 min. Cross-sections were then dehydrated in an ascending ethanol dilution series (70 %, 80 %, 90 %, 95 %, 100 % (2x)) and cleared in xylol twice for 2 min. Paraffin sections were then mounted with the xylene-based DPX mounting media for histology (Sigma-Aldrich). H&E stained sections were stored under the fume hood for 24 h before imaging by bright-field microscopy using the Zeiss Axio Lab.A1 microscope (Carl Zeiss).

### Immunofluorescence staining of testes cryosections

PFA-fixed murine testis cryosections stored at -20°C were defrosted and dried at room temperature for 15 min before their rehydration by washing in PBS and rinsing in distilled water. After drying the slides at RT for 10-15 min, cryosections were blocked (PBS supplemented with 1 % BSA, 2 % donkey serum and 0.3 % Triton X-100) for 1 h at RT while covered with parafilm to prevent evaporation. Primary antibodies diluted in PBST (0.3 % Triton X-100/PBS) supplemented with 1 % BSA were added and incubated at 4 °C overnight. On the next day, sections were washed three times for 5 min with PBST, and fluorescently-conjugated secondary antibodies diluted in PBST supplemented with 1 % BSA, were added and incubated in the dark for 1 h at RT. The sections were then washed three times for 5 min with PBST and slides were mounted with Fluoromount-G containing DAPI and photobleaching inhibitors (SouthernBiotech). All steps were performed in a dark humid chamber to prevent evaporation and photobleaching. Stained sections were stored at 4 °C for 24 h prior to imaging by immunofluorescence microscopy. Images were taken using the Zeiss Observer.Z1 epifluorescence microscope (Carl Zeiss) and the ZEN 2012 (blue edition) software (Carl Zeiss). Antibodies used are listed in Suppl. Table 3.

### RNA extraction and qRT-PCR quantification

RNA was extracted from cell pellets using the Trizol-containing reagent peqGold TriFAST according to the manufacturer’s instructions (PeqLab). RNA pellets were resuspended in RNase-free water, and DNA digestion was performed prior to RNA quantification. 0.5–1 µg of RNA was reverse transcribed to cDNA using the High Capacity cDNA Reverse Transcription Kit (Applied Biosystems) according to the manufacturer’s instructions. The cDNA was then diluted 1:5, and a relative quantification of specific genes was performed in a Bio-Rad qCycler using either TaqMan probes in iTaq Universal Probes Supermix or specific primer pairs in iTaq Universal SYBR Green Supermix (BioRad). Probes and Primers used are listed in Suppl. Table 2.

### Protein extraction and Western blotting

Cell pellets were lyzed in RIPA buffer (20 mM Tris–HCl pH 7.5, 150 mM NaCl, 1 mM Na_2_EDTA, 1 mM EGTA, 1 % NP-40, 1 mM Na_3_VO_4_, 1 % sodium deoxycholate, 2.5 mM sodium pyrophosphate, 1 mM glycerophosphate) supplemented with protease inhibitors and protein lysates were pre-cleared by centrifugation and quantified using the BCA assay kit (Pierce) according to the manufacturer’s instructions. Protein lysates were then denatured by incubation with SDS buffer (12 % glycerol, 60 mM Na_2_EDTA pH 8, 0.6 % SDS, 0.003 % bromophenol blue) for 10 min at 95 °C and separated in SDS-PAGE gels in Laemmli buffer (25 mM Tris, 192 mM glycine, 0.1 % SDS). Proteins were then wet transferred to a nitrocellulose membrane in transfer buffer (25 mM Tris–HCl pH 7.6, 192 mM glycine, 20% methanol, 0.03% SDS) and membranes were blocked with 3 % BSA in 1× TBST (50 mM Tris–HCl pH 7.6, 150 mM NaCl, 0.05 % Tween-20) prior to overnight incubation at 4 °C with the required primary antibodies. After washing the membrane three times with 1× TBST, they were incubated with a suitable HRP-coupled secondary antibody for 1 h at RT, followed by three washing steps with 1×TBST. Membranes were developed with the ECL substrate kit (Pierce) according to the manufacturer’s instructions. Antibodies used are listed in Suppl. Table 3.

### Cell culture

Derivation of wild type murine ESCs (WT, *Trim71*^*fl/fl*^) from conditional *Trim71* full knockout mice (*Trim71*^*fl/fl*^; *Rosa26-CreERT2*) was previously described^28^. *Trim71* knockout murine ESCs (KO, *Trim71*^*-/-*^) were generated from wild type ESCs by addition of 500 nM of 4-hydroxytamoxifen in their culture media for 48 h, followed by further culture for 72 h to achieve full protein depletion. ESCs were cultured in 0.1 % gelatin-coated dishes and maintained in 2i+LIF media (DMEM knockout media supplemented with 15 % FCS, 1 % penicillin-streptomycin, 0.1 mM NEAA, 2 mM L-GlutaMAX, 100 µM β-mercaptoethanol, 0.2 % in-house produced LIF, 1 µM of MEK/ERK inhibitor PD0325091 and 3 µM of GSK-3 inhibitor CHIR99021).

The human hepatocellular carcinoma cell line HepG2 and the human embryonic carcinoma cell lines JKT-1, NCCIT, NTERA-2, TCam-2 and 2102EP were acquired from ATCC. HepG2, NCCIT and TCam-2 were cultured in RPMI 1640 media supplemented with 10% FBS and 1% penicillin–streptomycin antibiotic solution. JKT-1, NTERA -2 and 2012EP cells were cultured in DMEM media supplemented with 10 % FBS and 1 % penicillin-streptomycin antibiotic solution.

### *In vitro* differentiation of ESCs into PGCLCs

ESCs growing under naїve pluripotency (d0) conditions (2i+LIF media on 0.1 % gelatine-coated dishes) were primed to EpiSCs (resembling post-implantation epiblast-derived stem cells) for two days by plating 10^5^ cells per well on 12-well plates coated with 20 µg/ml fibronectin in priming media (N2B27 media supplemented with 1 % penicillin-streptomycin, 2 mM L-GlutaMAX, 1 % KSR, 20 ng/ml Activin A and 12 ng/ml bFGF). After 24 h (d1), fresh priming media was provided. After 48 h (d2), EpiSCs were detached with Accutase Stem Cell Pro and 5000 cells per well were plated on a suspension U-bottom 96-well dish in specification media (GMEM media supplemented with 15 % KSR, 1 % penicillin-streptomycin, 0.1 mM NEAA, 2 mM L-GlutaMAX, 100 µM β-mercaptoethanol, 1 mM sodium pyruvate, 0.2 % in-house produced LIF, 50 ng/ml EGF, 100 ng/ml SCF and 500 ng/ml BMP4). After 6 days in this media (d8), cells growing in spheroids were recovered, detached by trypsinization for 10 min at 37°C and either used for RNA extraction or double-stained with anti-ITGB3/CD61 and anti-SSEA-1/CD15 antibodies to determine the number of PGCLCs by flow cytometry. For staining, trypsinized cells were washed in 0.1 % BSA/PBS, incubated with the antibodies diluted in 0.1 % BSA/PBS for 15 min at 4°C in the dark and washed again in 0.1 % BSA/PBS. Isotype control antibodies were used to control single stainings (data not shown). These protocols were adapted from a previous work^46^. Antibodies used are listed in Suppl. Table 3.

### Generation of NCCIT cells with *TRIM71* frameshift mutations

Gene editing using the CRISPR/Cas9 system was carried out using the plasmid pSpCas9(BB)-2A-GFP (PX458), which was a kind gift from Feng Zhang (Addgene plasmid #48138), after the insertion of specific sgRNAs targeting TRIM71 RING (sgRNA △RING 5′-CACCGCTCGCAGACGCTCACGCTGT-3′) or NHL (sgRNA △NHL6 5′-CACCGCACAACGATCATTCCGCTGG-3′) domains. NCCIT cells were transfected with PX458 empty vector (EV), TRIM71 ΔRING vector or TRIM71 ΔNHL6 vector using Lipofectamine Stem Transfection Reagent (Invitrogen) following the manufacturer’s instructions. 48 h post-transfection, transfected (GFP-positive) cells were sorted by FACS and plated as a bulk population. Growth competition assays (see below) were performed with sorted bulk populations. Generation of single clones from the bulk populations was also performed for subsequent Sanger sequencing and Western blot analysis.

### Growth competition assays and NGS allele frequency analysis via Illumina MiSeq

In order to analyze the growth behavior of CRISPR/Cas9-edited NCCIT cells, GFP-positive sorted bulk EV-transfected cells (WT) were mixed 1:1 with either TRIM71 sgRNA ΔRING-transfected cells or TRIM71 sgRNA ΔNHL6-transfected cells (d0) and cultured (2×10^5^ cells per well in a 6-well plate) for three weeks to investigate the population dynamics. Samples (2×10^5^ cells) were taken at d0, d3, d7, d14 and d21 for genomic DNA extraction. A double PCR strategy was then applied for the preparation of barcoded amplicons. In a first PCR reaction, the region around the CRISPR/Cas9-targeted site was amplified using site-specific forward and reverse primers with a common 5′ overhang. After PCR product cleaning, a second PCR was conducted with Illumina index primers which bind to the common 5′overhangs of the first primers and also contain unique barcode/index sequences followed by specific 5′ (P5) or 3′ (P7) adaptor sequences. PCR products were then gel-purified and sequenced on an Illumina MiSeq sequencing platform via amplification with common primers binding to P5/P7 adaptor sequences. Data analysis was performed using CRISPResso2^75^ with the quality cut-off set at 30 and the minimum identity score for the alignment being adjusted to 50. After analysis of the sequencing data, single reads were categorized as wild type, frameshift mutations and in-frame mutations, and allele frequencies were calculated and displayed as percentages in a pie chart. Comparing changes in the distribution of single reads for wild type (EV) and frameshift mutants TRIM71 ΔRING or TRIM71 ΔNHL6 over time gives insights into the dynamics of each cell populations within the initially 1:1 mixed culture. Allele frequencies of pure wild type (EV) were also analyzed at d0 and d21 as a control. Analysis of allele frequencies of pure WT, TRIM71 ΔRING and TRIM71 ΔNHL6 populations before (d0 pure) and after mixing them 1:1 (d0 1:1) ensured that no PCR product was biased over another based on theoretically expected vs. observed/measured allele frequencies.

### Exome sequencing study population

The study population consisted of 1025 individuals (MERGE cohort), of whom 908 otherwise healthy men presented with quantitative spermatogenic impairment. Patients attended the Centre of Reproductive Medicine and Andrology (CeRA) at the University Hospital Münster or the Clinic for Urology, Pediatric Urology and Andrology in Gießen. A subgroup of 247 selected patients was used to investigate novel genetic causes of the SCO phenotype (SCO subcohort). They had previously undergone testicular biopsy and presented with complete and bilateral absence of germ cells. In addition to this subgroup, 89 male subjects with complete spermatogenesis also pertaining to the MERGE cohort were included in the study as controls. All individuals gave written informed consent and the study protocol has been approved by the appropriate ethics committees according to the Declaration of Helsinki (Münster: Kennzeichen 2010-578-f-S, Gießen: No. 26/11). As further controls, we utilized a Dutch cohort of 5784 proven fathers sequenced at the Radboudumc genome diagnostics centre in Nijmegen, the Netherlands. These were healthy fathers of a child with severe developmental delay and all parents underwent routine exome sequencing for trio analysis. The fertility of the fathers is expected to be similar to an unselected sample of the population.

### Exome sequencing data generation and analysis

In order to perform exome sequencing, genomic DNA was isolated from the subjects’ peripheral blood applying standard procedures. Target enrichment, library capture, exome sequencing and variant calling were performed as previously described^64^. Briefly, DNA was enriched according to Agilent’s SureSelect^QXT^ Target Enrichment for Illumina Multiplexed Sequencing Featuring Transposase-Based Library Prep Technology or Twist Bioscience’s Twist Human Core Exome protocol. For library capture, SureSelectXT Human All Exon Kits (Agilent) or Human Core Exome (Twist Bioscience) were used. Sequencing was then conducted on Illumina HiScan®SQ, Illumina NextSeq®500/550 or Illumina HiSeqX® systems.

Exome data analysis was conducted on 247 SCO patients using our platform Sciobase. We filtered genes based on the predicted effect of the variant on translated protein (including only stop-gain, frameshift, and splice donor/acceptor variants) and the allele frequency in the general population (including only rare variants (MAF ≤ 0.01) as listed in gnomAD genomic database (v2.1.1: https://gnomad.broadinstitute.org)). This first filtering step resulted in the selection of 3997 genes. We then developed an application termed Haystack to facilitate the search for previously unknown causes of genetic diseases, technically applicable to exome data from patients with any clinical phenotype. In brief, Haystack aggregates information from multiple databases in an R Shiny application and allows for filtering based on the aggregated data, thus minimizing the manual effort required for candidate gene selection. All source code is publicly available at GitHub (https://github.com/MarWoes/haystack). We used Haystack to filter variants from our SCO patients for quality (read depth ≥10, variant depth ≥4 and frequency ≥40%). The remaining 2870 genes were filtered again based on genotype and allele frequency in gnomAD (MAF≤ 0.01 for homozygous variants and MAF ≤ 0.001 for heterozygous and X-linked variants) as well as on expression levels in the testes (including only genes whose expression in testis was above 50% of its maximum expression across all tissues according to the GTEx portal (v7: https://gtexportal.org)). Last, genes with LoF variants present in individuals with intact spermatogenesis were excluded from the selected 949 genes, leaving a total of 721 genes with LoF variants specifically found in SCO patients. *TRIM71* was found among these genes in a patient with an SCO phenotype (subject M364). This individual’s DNA was subjected to Sanger sequencing according to standard procedures in order to confirm his *TRIM71* frameshift variant. PCR was performed using the primers: F: 5′-TGCAGGCCTAATCGATGCAT-3′; R: 5′-GAAGAAGCTGCGTTGCCCTC-3′.

For the statistical analysis of exome data, the number of alleles affected and unaffected by LoF variants in *TRIM71* in all gnomAD subjects was compared to affected and unaffected alleles in the 908 patients with quantitative spermatogenic impairment (MERGE cohort) and the 247 SCO patients (SCO subcohort). A two-sided Fisher’s exact test was performed using the Woolf logit method.

In a subsequent step, exome data of the entire MERGE cohort and the Dutch father cohort was screened for rare (see MAF cutoffs above) missense variants in *TRIM71* (NM_001039111). The Z score for *TRIM71* was obtained from gnomAD database. Predicted pathogenicity was assessed using the scoring algorithms CADD (v1: https://cadd.gs.washington.edu/), PolyPhen (v2: http://genetics.bwh.harvard.edu/pph2), SIFT (v2: https://sift.bii.a-star.edu.sg), and Mutation Taster (v6: http://www.mutationtaster.org). Known genetic causes of azoospermia, such as chromosomal aberrations and AZF deletions were excluded in all infertile patients with LoF or missense variants in *TRIM71*. Furthermore, all patients carrying LoF or missense variants in *TRIM71* (listed in Table 1) were evaluated for relevant variants in other genes (n=181, Suppl. Table 1) that could explain the patients’ infertility. The majority of examined genes (n=170, Suppl. Table 1A) were taken from a review on the genetics of male infertility^76^. The genes were selected based on associated phenotypes and had been rated with limited to definitive clinical evidence. The list was amended by adding 11 recently published genes (Suppl. Table 1B). Specifically, the patients’ exome data was screened for stop-gain, frameshift, and splice acceptor/donor variants as well as missense variants with a CADD score ≥ 20. Only rare variants (gnomAD MAF ≤ 0.01 for homozygous and MAF ≤ 0.001 for heterozygous variants) were taken into account. For recessive genes, variants were only considered if found in homozygous or likely compound-heterozygous state (two variants in the same gene in the same patient).

## Acknowledgements

We thank all members of the Kolanus lab and the LIMES institute for general advice and discussion. LTF held a stipend from Bayer AG. WK is funded by the Deutsche Forschungsgemeinschaft (DFG, German Research Foundation) under Germany’s Excellence Strategy EXC2151 – 390873048. We are indebted to all subjects having donated their DNA for research use in our studies. The technical work of Christina Burhöi and Judith Ryll is gratefully acknowledged. This work was carried out within the frame of the German Research Foundation (DFG) funded Clinical Research Unit ‘Male Germ Cells: from Genes to Function’ (CRU326, grants to NN, CF and FT).

## Author contributions

LTF, JE, YP and SM designed and performed experiments. LTF and JE wrote the manuscript. SM and SS developed the germline-specific *Trim71* KO mouse. YP, SM, SS and LTF performed *in vivo* and *in vitro* experiments. MH and YP performed NGS Illumina MiSeq. NN and SDP provided testis single-cell RNAseq data. SK and DF provided samples and histological evaluation for patients from Münster and Gießen, respectively. JE and MW developed the variant filtering tool Haystack. JE performed bioinformatics analyses. MO provided genetic data for the cohort of Dutch fathers. WK, FT, HS and CF supervised experimental design, data analysis and manuscript writing.

## Notes

### Competing Interest Statement

The authors have declared no competing interest.

