## Supplementary figures and tables for "*TRIM71* deficiency causes germ cell loss during mouse embryogenesis and promotes human male infertility"

### Supplementary Figure 1

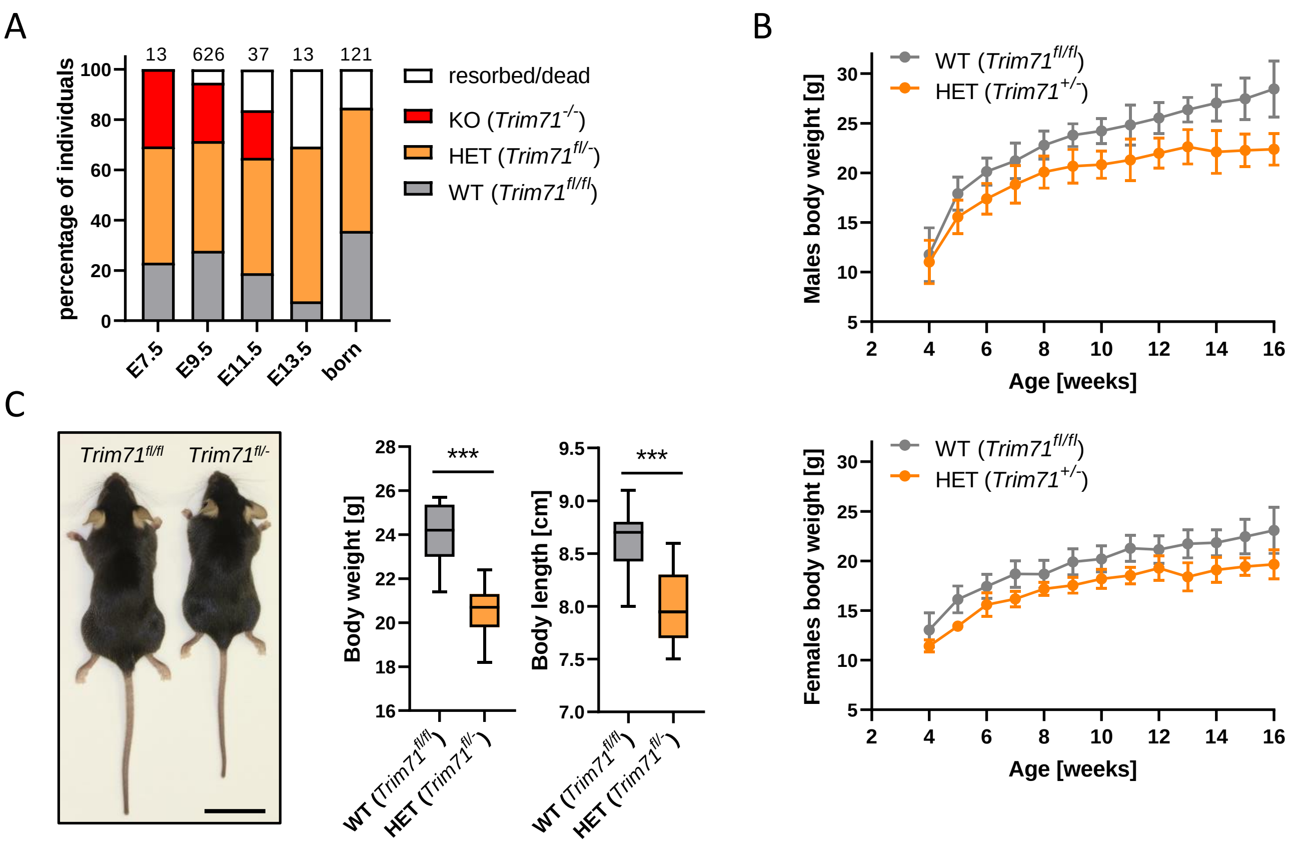

**Supplementary Figure 1 (relative to Figure 1). Heterozygous *Trim71* animals reveal a haploinsufficiency of *Trim71* gene. A)** Genotyping statistics of embryos from *Trim71* heterozygous intercrosses depicted as percentages. Embryonic lethality is apparent in by an alteration of Mendelian ratios in the neonatal (born, n=121) population (***P-value < 0.005, Chi square test), with embryos resorbed between E9.5-13.5. The numbers of analyzed individuals in each case are indicated above the columns. **B)** Growth curves (weight gain over time) for *Trim71* wild type (WT, *Trim71*$^{fl/fl}$) versus *Trim71* heterozygous (HET, *Trim71*$^{fl/-}$) mice for male (top graph) and female (bottom graph) mice. Error bars represent SEM (n=3-15). Weight gain overtime was significantly different between wild type and heterozygous mice for both males and females: ***P-value<0.005 (unpaired Student's t-test on area under curve (AUC) measurements). **C)** Representative image of adult male wild type and *Trim71* heterozygous mice accompanied by a quantification of body weight [g] and body length [cm] of adult (10-week-old) males. Scale bar represents 2 cm. Graphs represent Tukey plots (n=11-15). ***P-value < 0.005 (unpaired Student's t-test).

### Supplementary Figure 2

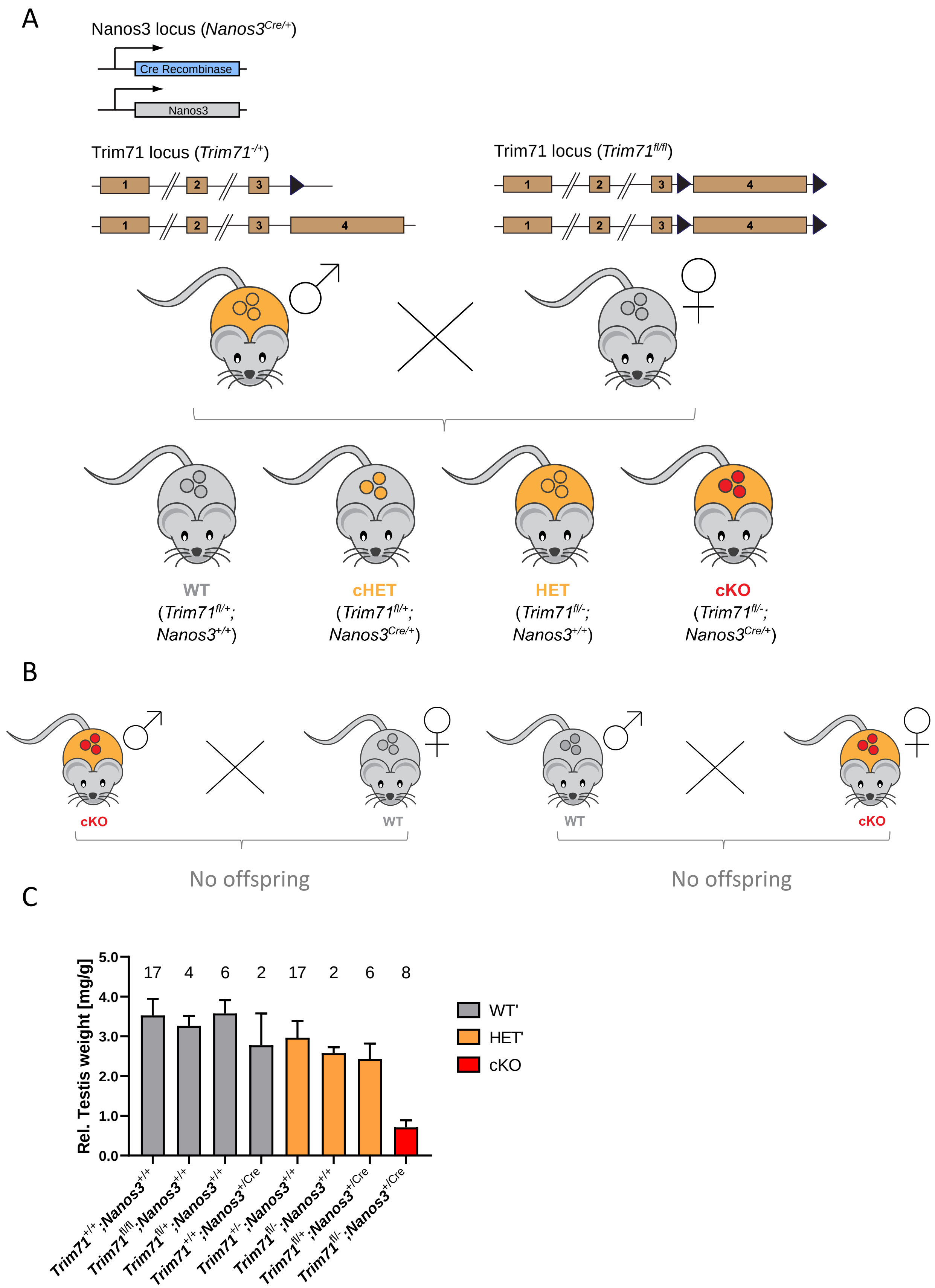

**Supplementary Figure 2 (relative to Figures 2 and 3). *Trim71* is essential for fertility. A)** Schematic representation for the generation of a germline-specific *Trim71* conditional knockout (cKO) mouse. Heterozygous (*Trim71*$^{+/-}$) males additionally carrying the *Nanos3*-Cre allele (*Nanos3*$^{Cre/+}$) were crossed with homozygously floxed *Trim71* (*Trim71*$^{fl/fl}$) females. The circles depicted inside the mice bodies represent the germline in adult males and females. Body and germline colors are indicative of the genotype: grey = WT, wild type (+/+, fl/fl or +/fl); orange = HET, heterozygous (+/- or fl/-) *Trim71* deletion; red = cKO, homozygous (-/-) *Trim71* deletion. **B)** Schematic representation of intercrosses for the functional characterization of *Trim71* cKO mice. No male or female germline-specific *Trim71* cKO mouse was able to generate offspring in the course of several months. **C)** Testis weight [mg] relative to total body weight [g] of adult male mice with different *Trim71* and *Nanos3* genotypes. In this case, colors are indicative of the *Trim71* genotype in the germline, regardless of *Nanos3* genotype. Error bars represent SEM and the numbers of analyzed individuals in each case are indicated above the columns.

### Supplementary Figure 3

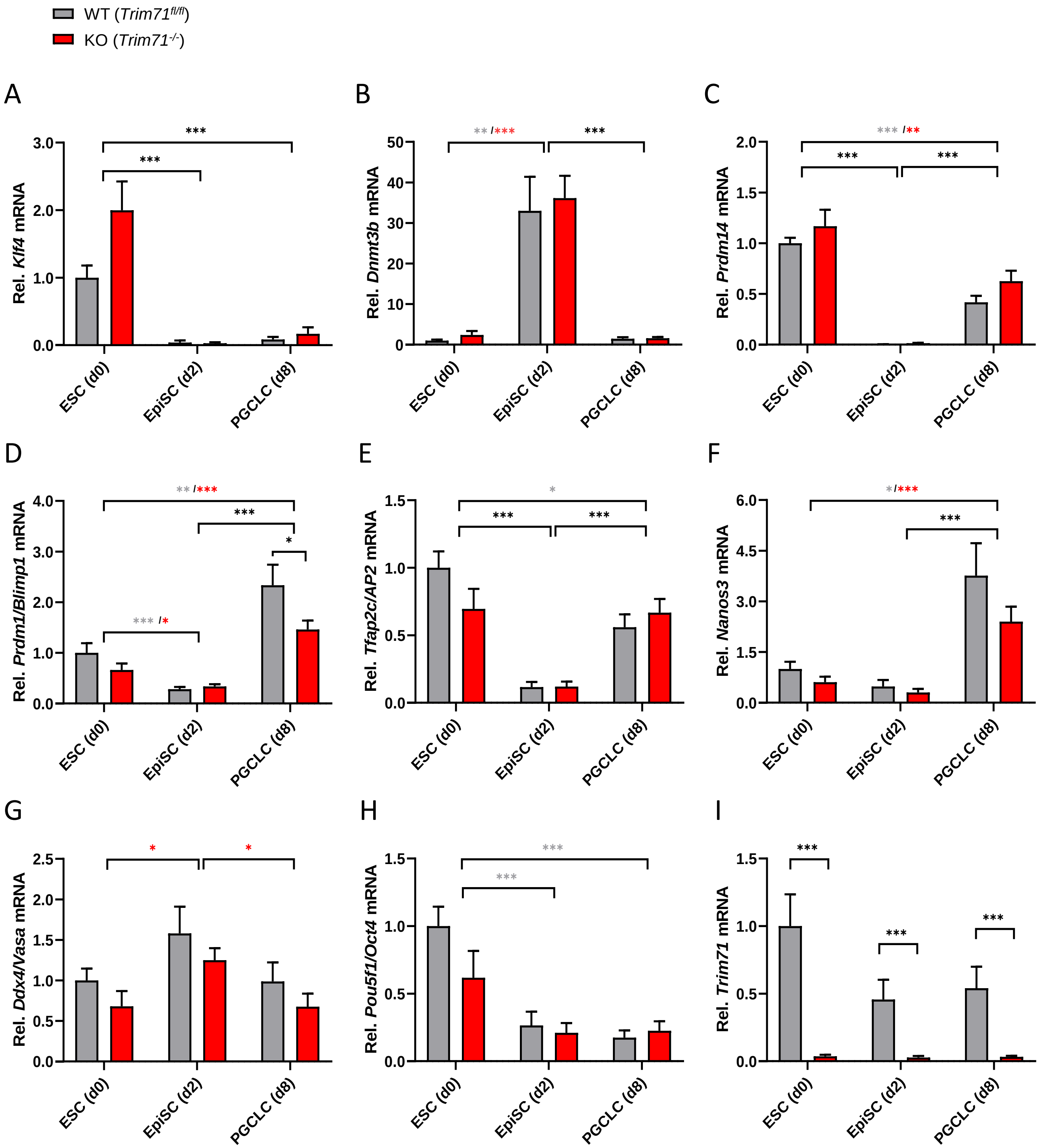

**Supplementary Figure 3 (relative to Figure 4). Monitoring *in vitro* differentiation of ESCs into PGCLCs.** qRT-PCR analysis of the indicated markers to monitor the priming and specification processes of mouse ESCs into PGCLCs, measured at d0 (ESCs, naïve pluripotency), d2 (EpiSCs, primed pluripotency) and d8 (PGCLCs, representing E7.5 specified pre-migratory PGCs). As detailed in the main text, *Klf4* is a marker of naïve pluripotency and *Dnmt3b* of primed pluripotency, while *Trim71* and *Pou5f1/Oct4* expression is expected in both naïve and primed stem cells. *Prdm14, Prdm1/Blimp1, Tfap2c* and *Nanos3* are early PGC markers expected to be upregulated at d8 compared to d2. *Ddx4/Vasa* is a late PGC marker expected to be upregulated after PGC migration to the genital ridges (E10.5). ***P-value < 0.005, **P-value < 0.01; P-value < 0.005; black stars represent equal statistical significance for both genotypes, while grey and red stars represent specific significance for wild type (WT, *Trim71*$^{fl/fl}$) and *Trim71* knockout (KO, *Trim71*$^{-/-}$) cells, respectively (unpaired Student's t-test).

### Supplementary Figure 4

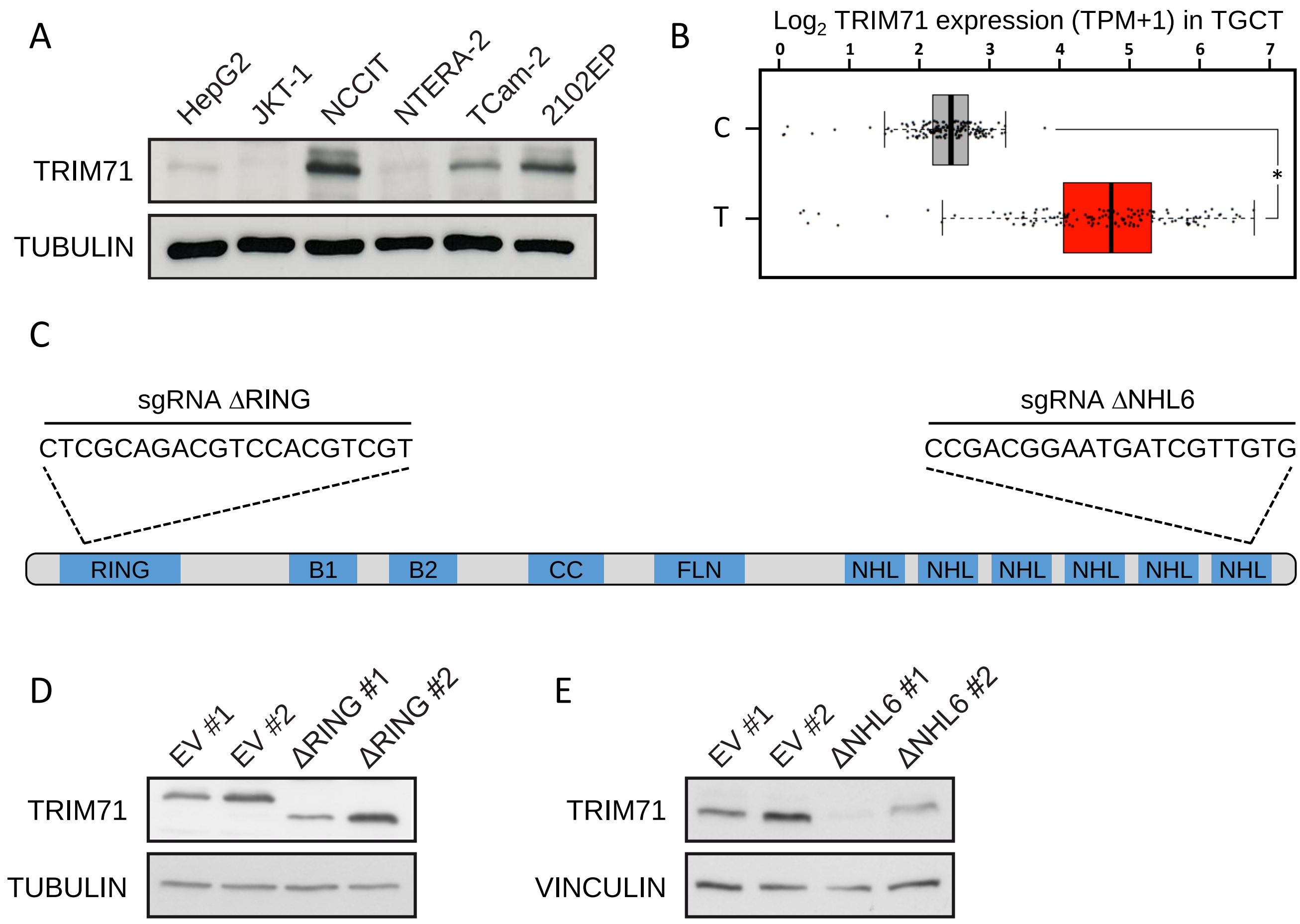

**Supplementary Figure 4 (relative to Figure 5). Generation of *TRIM71* mutations via CRISPR/Cas9 in NCCIT cells. A)** Western blot showing TRIM71 protein expression in several germ cell tumor (GCT)-derived embryonic carcinoma cell lines. TUBULIN was used as a loading control. 20 µg of protein were loaded per lane. **B)** *TRIM71* expression is upregulated in human testicular germ cell tumor (TGCT) patients. Box plot graph was obtained from GEPIA (http://gepia2.cancer-pku.cn) and shows *TRIM71* mRNA levels in human control (C, n=165) and tumor (T, n=137) samples. TPM = transcript counts per million. **C)** Schematic representation of TRIM71 structural domain organization indicating the location and target sequences (single guide RNA = sgRNA) for the generation of NCCIT cells with *TRIM71* frameshift mutations in the RING domain (ΔRING) and the NHL domain (ΔNHL6) via CRISPR/Cas9. **D)** Western blot showing TRIM71 protein in two single NCCIT clones targeted with sgRNA ΔRING including wild type cells transfected with an empty vector (EV). Of note, TRIM71 RING mutant NCCIT cells (ΔRING) express a RINGless protein of 83 kDa due to the usage of an alternative in-frame ATG codon located downstream of the targeted sequence. **E)** Western blot showing TRIM71 protein in two single NCCIT clones targeted with sgRNA ΔNHL6 including wild type cells transfected with an empty vector (EV). For TRIM71 NHL mutant NCCIT cells (ΔNHL6) the deletion of the last C-terminal 24 aa (clone #1) renders an unstable protein which seems to be degraded while the deletion of the last C-terminal 18 aa (clone #2) renders a stable C-terminal truncated protein of 92 kDa.

### Supplementary Figure 5

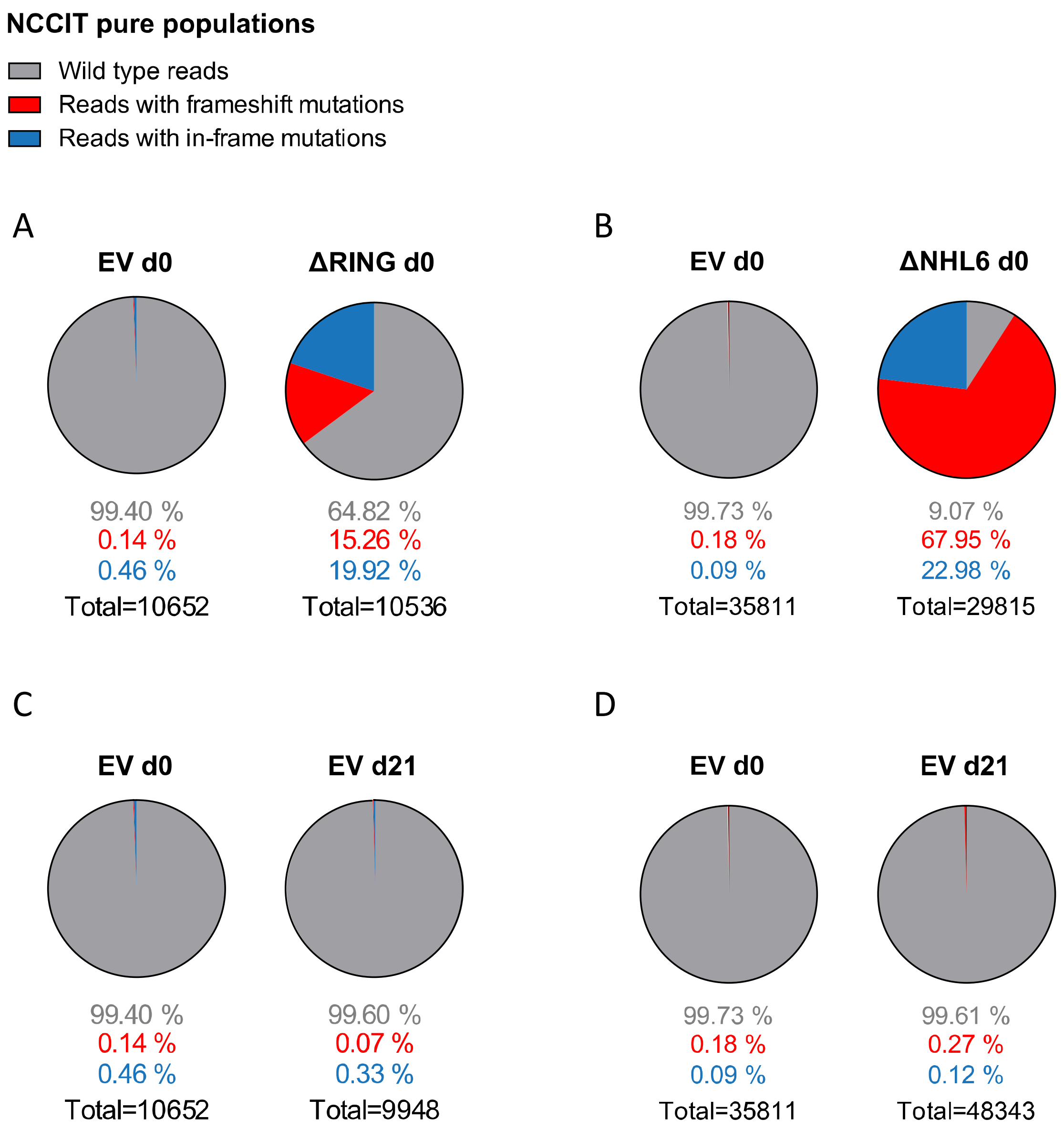

**Supplementary Figure 5 (relative to Figure 5). NGS analysis via Illumina MiSeq platform of pure NCCIT wild type and *TRIM71* mutant populations. A)** Pie charts showing allele frequencies at d0 for wild type NCCIT cells transfected with an empty vector (EV) or with a sgRNA targeting TRIM71's RING domain (ΔRING). The depicted pure populations were then mixed 1:1 and cultured for growth competition assays in which allele frequencies were measured at different time points via Illumina Miseq analysis (corresponding to Fig. 5A). **B)** Pie charts showing allele frequencies at d0 for wild type NCCIT cells transfected with an empty vector (EV) or with a sgRNA targeting TRIM71's NHL domain (ΔNHL6). The depicted pure populations were then mixed 1:1 and cultured for growth competition assays in which allele frequencies were measured at different time points via Illumina Miseq analysis (corresponding to Fig. 5B). **C)** Pie charts showing allele frequencies at d0 and d21 for wild type NCCIT cells transfected with an empty vector (EV) from A. **E)** Pie charts showing allele frequencies at d0 and d21 for NCCIT cells transfected with an empty vector (EV) from B.

### Supplementary Figure 6

A

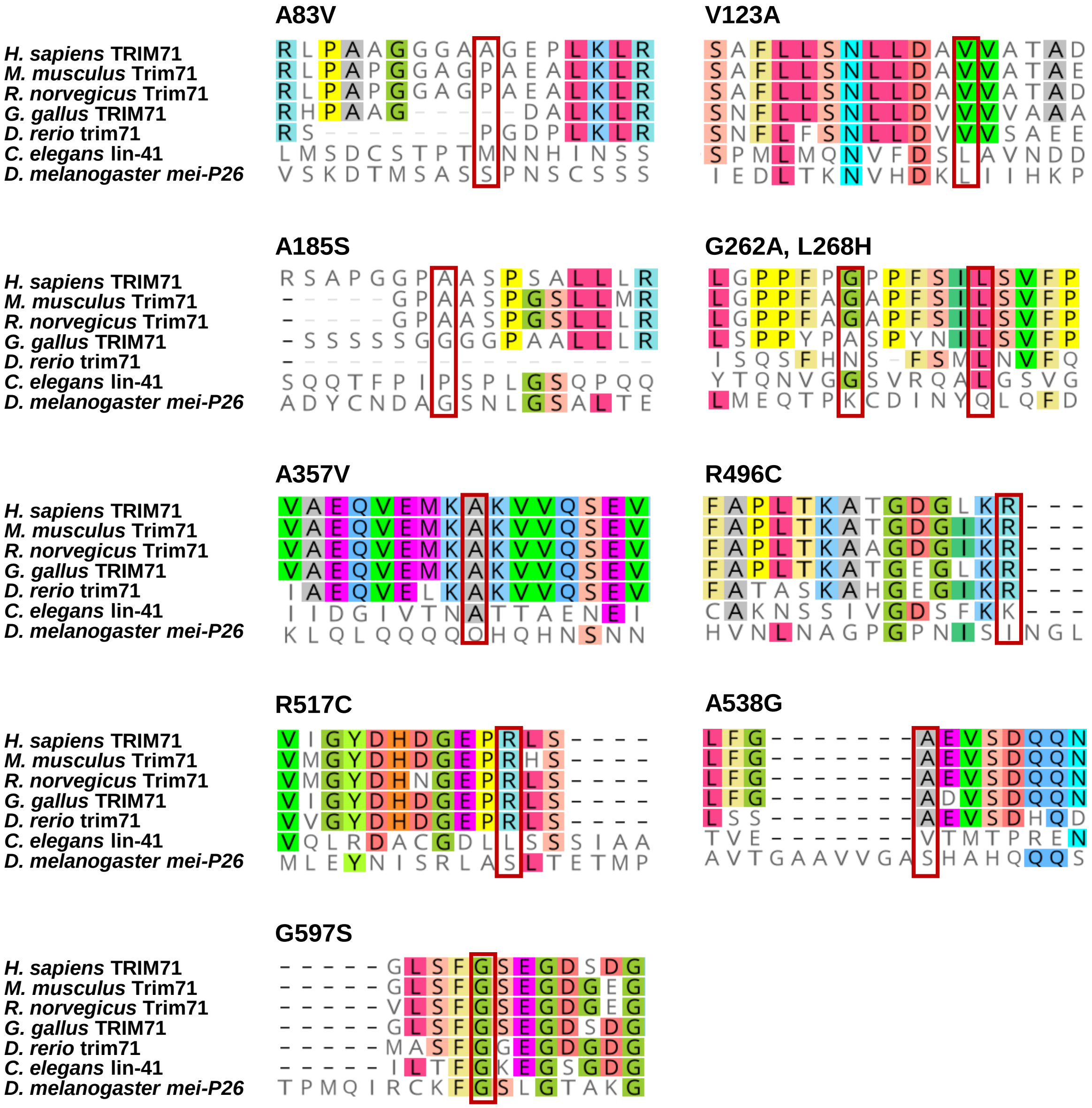


B

***TRIM71* LoF variant c224_240dup/p.(Gly81CysfsTer24)**

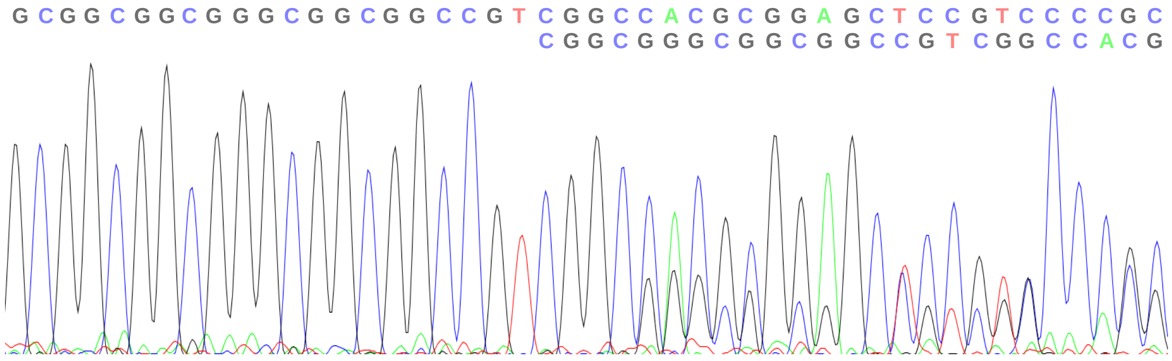

**Supplementary Figure 6 (relative to Figure 6). Analysis of the evolutionary conservation of *TRIM71* genetic variants found in male infertile patients. A)** Sequence conservation logos for the *TRIM71* genetic variants found in individuals of the MERGE cohort. Sequence logos were obtained from MEGA/ClustalW and Geneious Prime. Affected amino acids are indicated in red boxes across all examined vertebrate and invertebrate species. **B)** Electropherogram resulting from Sanger sequencing of DNA from an SCO patient (subject M364) showing the duplication c.224_240dup resulting in the frameshift p.(Gly81CysfsTer24) in *TRIM71* (reverse strand).

### Supplementary Table 1

| **Supplementary Table 1A. Infertility-associated genes (Reviewed in Oud *et al.,* 2019. Human Reproduction)** | | | | | | | | |
|---|---|---|---|---|---|---|---|---|
| ABCA1 | CCDC141 | DMRT1 | FGFR1 | INSL3 | NOS1 | REC8 | SPO11 | UBE2B |
| ADGRG2 | CCDC155 | DNAAF2 | FLNA | JAG1 | NOTCH1 | RELN | SRA1 | UBR2 |
| AK7 | CCDC39 | DNAAF4 | FSHB | KDM3A | NR0B1 | RNF220 | SRD5A2 | USP26 |
| AKAP4 | CCDC40 | DNAAF5 | FSHR | KISS1R | NR5A1 | RSPH1 | SRY | VAMP7 |
| AMH | CDC14A | DNAH1 | FSIP2 | KLHL10 | NRAS | RSPH3 | STK36 | WDR11 |
| AMHR2 | CEP135 | DNAH9 | GALNTL5 | LHB | NSMF | RSPH9 | STX2 | WDR66 |
| ANOS1 | CEP290 | DNAI1 | GAS8 | LHCGR | PANK2 | RSPO1 | SUN5 | WT1 |
| APOA1 | CFAP43 | DNAI2 | GATA4 | LRRC6 | PDHA2 | SECISBP2 | SYCE1 | XRCC2 |
| AR | CFAP44 | DNAJB13 | GH1 | MAGEB4 | PIH1D3 | SEMA3A | SYCP3 | ZMYND15 |
| AURKC | CFAP69 | DNMT1 | GNRH1 | MAMLD1 | PKD1 | SLC26A3 | TAC3 | ZPBP |
| BMP4 | CFTR | DNMT3B | GNRHR | MAP2K2 | PLCZ1 | SLIT2 | TACR3 | |
| BMP7 | CHD7 | DPY19L2 | GTF2H3 | MAP3K1 | PLEKHA5 | SOS1 | TAF4B | |
| BNC2 | CYP11A1 | E2F1 | HAUS7 | MC4R | PLK4 | SOX10 | TDRD6 | |
| BRAF | CYP11B1 | ERBB4 | HESX1 | MEI1 | PLXNA1 | SOX2 | TDRD7 | |
| BRDT | CYP17A1 | FANCA | HS6ST1 | MEIOB | PMFBP1 | SOX3 | TDRD9 | |
| BSCL2 | CYP19A1 | FANCM | HSD17B3 | MNS1 | POLR3B | SOX8 | TEX11 | |
| C11orf70 | CYP21A2 | FATE1 | HSD3B2 | MTOR | PROK2 | SOX9 | TEX14 | |
| CATSPER1 | DCC | FEZF1 | HSF2 | NANOS2 | PROKR2 | SPAG17 | TEX15 | |
| CATSPERE | DHH | FGF17 | HYDIN | NLRP3 | PSMC3IP | SPATA16 | TRIM37 | |
| CCDC103 | DMC1 | FGF8 | IL17RD | NNT | RBMXL2 | SPINK2 | TTLL5 | |

| **Supplementary Table 1B. Infertility-associated genes (reported recently in independent studies)** | |
|---|---|
| ADAD2 | Krausz et al., 2020. Genetics in Medicine |
| M1AP | Wyrwoll et al., 2020. AJHG |
| MSH4 | Krausz et al., 2020. Genetics in Medicine |
| RAD21L1 | Krausz et al., 2020. Genetics in Medicine |
| RNF212 | Riera-Escamilla et al., 2019. Human Reproduction |
| SHOC1 | Krausz et al., 2020. Genetics in Medicine |
| STAG3 | Riera-Escamilla et al., 2019. Human Reproduction. Van der Bijl et al., 2019. Human Reproduction |
| SYCP2 | Schilit et al., 2020. AJHG |
| TERB1 | Krausz et al., 2020. Genetics in Medicine & Salas-Huetos et al., 2020. Human Genetics |
| TERB2 | Salas-Huetos et al., 2020. Human Genetics |
| MAJIN | Salas-Huetos et al., 2020. Human Genetics |

**Supplementary Table 1C. Variants identified in infertility-associated genes in patients with TRIM71 variants**

| **Subject ID** | ***TRIM71* variant** | **Infertility-associated gene** | **Variant** |
|---|---|---|---|
| M1686 | c.1070C>T/p.(Ala357Val) | *SYCP2* | heterozygous, frameshift |
| M1083 | c.1486C>T/p.(Arg496Cys) | *NNT** | heterozygous, missense |
| M468 | c.1549C>T/p.(Arg517Cys) | *TERB1** | homozygous, missense |
| M468 | c.1549C>T/p.(Arg517Cys) | NOTCH1* | heterozygous, missense |
| M468 | c.1549C>T/p.(Arg517Cys) | TDRD7* | heterozygous, missense |
| M754 | c.1789G>A/p.(Gly597Ser) | *HSD3B2** | heterozygous, missense |
| M2141 | c.1789G>A(p.(Gly597Ser) | *CCDC141** | heterozygous, missense |

**Supplementary Table 1 (relative to Figure 6).** Analysis of exome data from infertile men carrying *TRIM71* variants for the identification of variants in other genes previously associated with male infertility in humans. *Variants of uncertain significance.

### Supplementary Table 2

| **Supplementary Table 2A. Genotyping primers** | |
|---|---|
| *Trim71_F1* | 5'-GAAAGGAGGCTAGCCAAAGG-3' |
| *Trim71_R1* | 5'-ATGCTGTACGGTAGGAGTCTTCC-3' |
| *Trim71_R2* | 5'-CACACAAAAAACCAACACACAG-3' |
| *Nanos3_F1* | 5'-CCAGCCATGGGGACTTTC-3' |
| *Nanos3_R1* | 5'-GGGACTGATAGATGGCAC-3' |
| *Nanos3_R2* | 5'-CAGAGGCCACTTGTGTAGCG-3' |
| **Supplementary Table 2B. qRT-PCR primers (SYBR Green)** | |
| *Klf4_F* | 5'-GCGAACTCACACAGGCGAGAAAC-3' |
| *Klf4_R* | 5'-TCGCTTCCTCTTCCTCCGACA-3' |
| *Prdm14_F* | 5'-CAGCGACTTCATTGCCAAAGGAG-3' |
| *Prdm14_R* | 5'-GCCGTCGATAAAATGGCTCAGG-3' |
| *Prdm1/Blimp1_F* | 5'-ACCCCTCATCGGTGAAGTCTACA-3' |
| *Prdm1/Blimp1_R* | 5'-CTCCTCTCTGGAATAGATCCGCCA-3' |
| *Ddx4/Vasa_F* | 5'-TCATACTTGCAGGACGAGATTTG-3' |
| *Ddx4/Vasa_R* | 5'-AACGACTGGCAGTTATTCCATC-3' |
| *18 sRNA_F* | 5'-GTAACCCGTTGAACCCCATTC-3' |
| *18 sRNA_R* | 5'-CCATCCAATCGGTAGTAGCGAC-3' |
| **Supplementary Table 2C. qRT-PCR probes (TaqMan)** | |
| *Trim71* | Mm01341471_m1 |
| *Sall4* | Mm00453037_s1 |
| *Lin28a* | Mm00524077_m1 |
| *Lin28b* | Mm01190673_m1 |
| *Pou5f1/Oct4* | Mm03053917_g1 |
| *Dmnt3b* | Mm01240113_m1 |
| *Tfap2c* | Mm00493473_m1 |
| *Nanos3* | Mm00808138_m1 |
| *Hprt* | Mm03024075_m1 |

**Supplementary Table 2 (relative to Methods).** Primers and Probes used for the amplification of murine genes in genotyping or qRT-PCR analyisis. F = forward; R = reverse. TaqMan probes were acquired from Applied Biosystems (Thermo Fischer Scientific).

### Supplementary Table 3

| Supplementary Table 3A. Primary Antibodies | | | | | |
|---|---|---|---|---|---|
| **Protein** | **Conjugate** | **Source Species** | **Dilution** | **Application** | **Company_Reference** |
| GCNA1/TRA98 | - | rat | 1:500 | IF | Abcam_ab82527 |
| WT1 | - | rabbit | 1:50 | IF | Abcam_ab89901 |
| TRIM71 | - | rabbit | 1:1000 | WB | Sigma_HPA038142 |
| TUBULIN | - | mouse | 1:2000 | WB | Sigma_T9026 |
| VINCULIN | - | mouse | 1:2000 | WB | Sigma_V9131 |
| THY1.2/CD90.2 | FITC | rat | 1:400 | FACS/MACS | Biolegend_140303 |
| SSEA-1/CD15 | Alexa 647 | mouse | 1:200 | FACS | Biolegend_125607 |
| ITGB3/CD61 | PE-Cy7 | armenian hamster | 1:400 | FACS | Biolegend_104317 |
| **Supplementary Table 3B. Secondary Antibodies** | | | | | |
| **Protein** | **Conjugate** | **Source Species** | **Dilution** | **Application** | **Company_Reference** |
| rat IgG | Alexa 488 | donkey | 1:250 | IF | Jackson ImmunoResearch_712-545-153 |
| rabbit IgG | Cy3 | donkey | 1:400 | IF | Jackson ImmunoResearch_711-165-152 |
| rabbit IgG | HRP | goat | 1:5000 | WB | Cell Signaling Technologies_7074 |
| mouse IgG | HRP | horse | 1:5000 | WB | Cell Signaling Technologies_7076 |

**Supplementary Table 3 (relative to Methods).** Antibodies used for immunofluorescence (IF), western blot (WB), flow cytometry (FACS) and magnetic-assisted cell sorting (MACS).
